# Disrupted endosomal trafficking of the Vangl-Celsr polarity complex underlies congenital anomalies in trachea-esophageal morphogenesis

**DOI:** 10.1101/2023.10.11.561909

**Authors:** Nicole A. Edwards, Scott A. Rankin, Adhish Kashyap, Alissa Warren, Zachary N. Agricola, Alan P. Kenny, Matthew J. Kofron, Yufeng Shen, Wendy K. Chung, Aaron M. Zorn

## Abstract

Disruptions in foregut morphogenesis can result in life-threatening conditions where the trachea and esophagus fail to separate properly, such as esophageal atresia (EA) and tracheoesophageal fistulas (TEF). The developmental basis of these congenital anomalies is poorly understood, but recent genome sequencing reveals that *de novo* variants in intracellular trafficking genes are enriched in EA/TEF patients. Here, we confirm that mutation of orthologous genes in *Xenopus* disrupts trachea-esophageal separation similar to EA/TEF patients. We show that the Rab11a recycling endosome pathway is required to localize Vangl-Celsr polarity complexes at the luminal cell surface where opposite sides of the foregut tube fuse. Partial loss of endosome trafficking or Vangl-Celsr complexes disrupts epithelial polarity and mutant cells accumulate at the fusion point, fail to downregulate Cadherin, and do not separate into distinct trachea and esophagus. These data provide insights into the mechanisms of congenital anomalies and general paradigms of tissue fusion during organogenesis.

## Introduction

Esophageal atresia (EA) and tracheoesophageal fistulas (TEF), which affect approximately 1 in 3000 newborns, are life-threatening congenital anomalies that arise when the embryonic foregut does not separate correctly into distinct trachea and esophageal tubes^1–4^. This results in esophageal discontinuity (EA) and/or a pathological connection between the trachea and esophagus (TEF). Rarer are laryngeal-tracheal-esophageal-clefts where, in some extreme cases, the foregut fails to separate along its entire length, essentially resulting in a complete TEF^5^. Roughly half of EA/TEF patients also have variably penetrant co-occurring anomalies in other organs, including gastrointestinal, pulmonary, cardiovascular, renal, and central nervous system, suggesting undefined syndromes^6,7^. The genetic etiology of tracheoesophageal anomalies is poorly understood and is associated with spontaneous *de novo* nucleotide variants, chromosomal rearrangements, copy number variation and indels in affected newborns^1,8^. Known variants only account for 10-15% of cases worldwide with the remaining having no identified risk gene. Even when the causative variants are known, why they result in tracheoesophageal anomalies is unclear since the cellular mechanisms governing foregut morphogenesis are poorly understood.

Tracheoesophageal morphogenesis begins when the foregut endoderm is patterned along the dorsal-ventral axis by an evolutionarily conserved Hedgehog-Wnt-BMP signaling network, resulting in dorsal Sox2+ esophageal progenitors and ventral Nkx2-1+ respiratory progenitors^9–12^. After patterning, the foregut constricts at the midline, and a population of epithelial cells on either side of the foregut tube that express both dorsal and ventral markers (Sox2/Nkx2-1/Islet1) come together and adhere, forming a transient epithelial bilayer^13,14^. Epithelial cells at the fusion point then progressively lose adhesion to one another and integrate into either the ventral or dorsal foregut as the surrounding Foxf1+ splanchnic mesoderm invades between the nascent trachea and esophagus^14^. Thus, the single foregut epithelial tube fuses, reorganizes its cellular junctions, and then undergoes fission with coordinated cell rearrangements to reconnect the epithelia into two separate tubes. How cell signaling and transcription factors regulate the cellular processes of tracheoesophageal morphogenesis is unclear.

Similar epithelial fusion and fission events are crucial for the morphogenesis of many tissues, including the heart, oral palate, eyes, inner ear, urogenital sinus, and neural tube; organ systems that can also be disrupted in complex EA/TEF patients, suggesting shared biological mechanisms^15^. Such tissue fusion and fission events involve dynamic and coordinated changes in cell adhesion and polarity^15,16^. For instance, studies of vertebrate neural tube closure and *Drosophila* retina morphogenesis indicate that the precise spatial and temporal coordination of cell adhesion and cell polarity across different epithelial populations is essential for proper tissue fusion. Less is known about how epithelial polarity and adhesion, likely downstream of Hedgehog and BMP signaling^17^, are regulated in tissue fission events like urogenital sinus-hindgut separation. A remaining challenge in the field is understanding how the proteins involved in cell adhesion, cell polarity, and cell junctions are orchestrated to regulate epithelial remodeling during morphogenesis^18–21^.

We recently found in *Xenopus* that Rab11+ recycling endosomes somehow regulate tracheoesophageal morphogenesis downstream of Hedgehog signaling^14^. Rab11 is a small GTPase required for the intracellular recycling of endosomal vesicles that cells use to transport newly synthesized proteins from the Golgi to the plasma membrane and to change the composition of specific plasma membrane domains by internalizing receptors and other cell surface proteins^22^. How recycling endosomes regulate epithelial remodeling in the foregut was unknown. However, recent whole-genome sequencing of EA/TEF patients who exhibit complex multi-organ phenotypes revealed an enrichment of potentially damaging *de novo* variants in genes involved in endocytosis and intracellular trafficking^23^. This finding and our previous observations prompted us to test the hypothesis that endosome trafficking represents a critical morphogenetic bottleneck in tracheoesophageal separation and to explore the underlying cellular mechanisms.

Here we demonstrate that the recycling endosome pathway is required upstream of the Vangl-Celsr cell polarity complex for proper fission of the foregut tube. Mechanistically, Rab11a localizes the transmembrane protein Vangl2 to the midline luminal cell surface where juxtaposing apical sides of the foregut touch, localize Cadherin, and adhere to one another to form the transient epithelial bilayer. Rab11a and Vangl-Celsr complexes at the contact surface appear to confer an “apical memory” essential for maintaining the cellular organization of transient bilayer and its subsequent resolution. A partial loss of 1) core endosome trafficking components, 2) orthologs of EA/TEF patient risk genes, or 3) the Vangl-Celsr proteins disrupt cell polarity and the orientation of cell division in the transient bilayer. Mutant cells accumulate in a disorganized septum, fail to internalize E-cadherin (Cdh1) from the rounding cell surfaces, and the foregut does not separate into trachea and esophagus, resulting in an extended tracheoesophageal fistula. This unexpected Vangl-Celsr activity is distinct from their well-known roles in planar cell polarity (PCP) which orients cells within the plane of the epithelium. Together, these results indicate that failure to maintain localized endosome-mediated epithelial polarity during foregut morphogenesis may be a common disease mechanism underlying EA/TEF in many patients and may have broad implications for tissue fusion-fission events in the morphogenesis of other organs.

## Results

### Mutation of endosome pathway genes implicated from EA/TEF patients cause TEF in Xenopus

Recent genome sequencing of complex EA/TEF patients revealed a significant enrichment of potentially damaging and missense *de novo* heterozygous variants in genes involved in intracellular trafficking and endocytosis^23^. This was intriguing considering our discovery that the recycling endosome pathway was somehow involved in tracheoesophageal morphogenesis downstream of Hedgehog signaling^14^. A meta-analysis of experimentally validated protein-protein interactions from the STRING database^24^ confirmed that many proteins encoded by patient genes interact with core recycling endosome components (Figure 1A).

**Figure 1.**
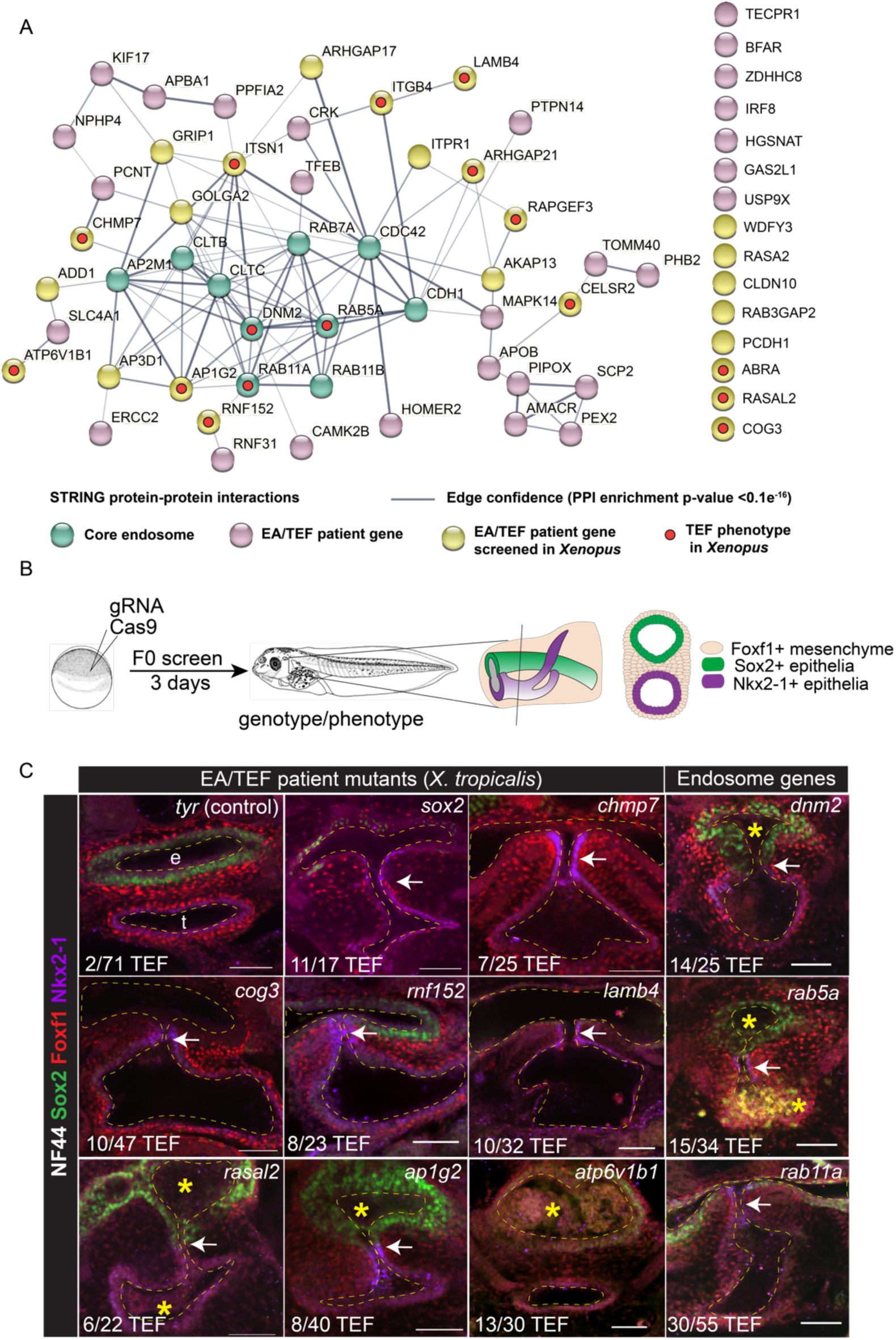
Mutating EA/TEF patient endosome trafficking genes causes TEDs in *Xenopus*. A) STRING Database interactome of endosome-related proteins with potentially pathogenic variants identified in and a curated list of core endosome pathway proteins and putative protein cargo. B) Experimental design of F0 CRISPR-Cas9 *X. tropicalis* mutagenesis screen to validate candidate risk genes. C) Confocal images of CRISPR-Cas9 *X. tropicalis* F0 mutants at NF44 stained for Sox2 (green), Foxf1 (red), and Nkx2-1 (purple). Hashed yellow lines indicate the trachea (t) and esophageal (e) lumens. Arrows indicate TEFs. Asterisks indicate dysmorphic or occluded esophagus or trachea. Numbers indicate the proportion of mutant tadpoles with TEF compared to the total mutants screened. Scale bars are 50 µm.

To systematically test the role of endosome trafficking in EA/TEF and determine if the patient variants were likely to be causative risk alleles, we mutated orthologous genes in *Xenopus tropicalis* using F0 CRISPR-Cas9 gene editing. We compared the resulting phenotypes to those caused by mutations in core endosome pathway genes. We designed guide RNAs (gRNAs) to create indel mutations early in the coding sequence that were predicted to disrupt protein function by creating a premature stop codon or missense mutations leading to nonsense-mediated decay^25,26^. Cas9-gRNA complexes were injected at the one-cell stage, and after three days of development, individual embryos were genotyped, and tracheoesophageal morphogenesis was assayed by immunostaining and confocal microscopy (Figure 1B). Guide RNAs that reproducibly caused EA/TEF-like phenotypes with a significant penetrance compared to controls would support the patient’s variant being involved in EA/TEF. In contrast, genes that did not reproducibly cause EA/TEF when mutated in *Xenopus* were likely not disease-causing in EA/TEF patients.

Confirming the result of 10 genes previously screened^23^ as well as testing 15 new genes, we found that mutations in 13 out of 25 orthologs of endosome-related patient risk genes result in significant variably penetrant tracheoesophageal anomalies in *Xenopus* (Figure 1C and Table S1; p<0.05). In each case, the foregut was correctly patterned into a ventral Nkx2-1+ tracheal domain and a dorsal Sox2+ esophageal domain with surrounding Foxf1+ mesenchyme, and the foregut underwent initial constriction at the midline. However, in many embryos, the foregut failed to separate into distinct tubes and exhibited a persistent fistula (Figure 1C, arrows). These phenotypes were similar to those caused by mutations in the known EA/TEF risk gene *sox2*^23^, and mutations in genes encoding key GTPases in endosome trafficking: *dynamin-2* (*dnm2)*, which regulates endocytosis at the cell surface, *rab5a*, which regulates early endosome vesicle trafficking, and *rab11a*^14^ that regulates recycling endosomes (Table S1). Most of the mutants that caused in TEF also exhibited a low penetrance (∼20%) of dysmorphic esophagus or trachea ranging from stenosis to a thick multilayered epithelium with multiple or occluded esophageal lumens full of cells (Figure 1C, asterisks; Table S1).

Genotyping demonstrated an 80% average indel mutation rate (% alleles mutated per embryo) across all gRNAs with a range of 58-90% (Table S1 and Figure S1A). Most gRNAs that reproducibly cause TEF had a mutation burden exceeding 75%. Sequencing analysis revealed that a given gRNA often produced a reproducible pattern of 1-3 different mutations (Figure S1B, source data in Table S2; see STAR methods for details on the genotype-phenotype analysis). Thus, while different cells in an embryo could have different indels, most cells likely had heterozygous or biallelic mutations. In cases when we had effective antibodies (Dnm2, Rab5a, and Rab11a), western blotting of foregut explants showed a dramatic loss of protein, and immunostaining of unilaterally injected embryos demonstrated that most of the cells (>90%) on the injected side had reduced target protein abundance (Figure S1C-D). Together, these data indicate that the F0 CRISPR strategy results in most embryos being more damaged than heterozygous but not complete nulls, with little cellular mosaicism.

Interestingly, in some mutants, we observed phenotypes in other tissues, including craniofacial, heart, eye, body axis, and gut coiling defects, organ systems that are also disrupted in many EA/TEF patients. However, we found no evidence of a generalized developmental delay based on developmental milestones such as melanocyte pigmentation and tail fin formation. As expected for genes in essential cellular pathways, some embryos with a very high mutation burden (>80%) exhibited gastrulation arrest and lethality before foregut morphogenesis.

Taken together, these results confirm that endosome trafficking is critical for tracheoesophageal morphogenesis in *Xenopus* and support the likelihood that variants in endosome-related genes may have caused EA/TEF in some patients.

### Dynamic endosome localization and epithelial remodeling during foregut morphogenesis

To better understand the cellular mechanisms by which endosome trafficking regulates *Xenopus* foregut morphogenesis, we performed a detailed immunostaining analysis of Dnm2, Rab5a, and Rab11a over 24 hours (at room temperature) when the foregut separates into distinct esophageal and tracheal tubes (Figure 2).

**Figure 2.**
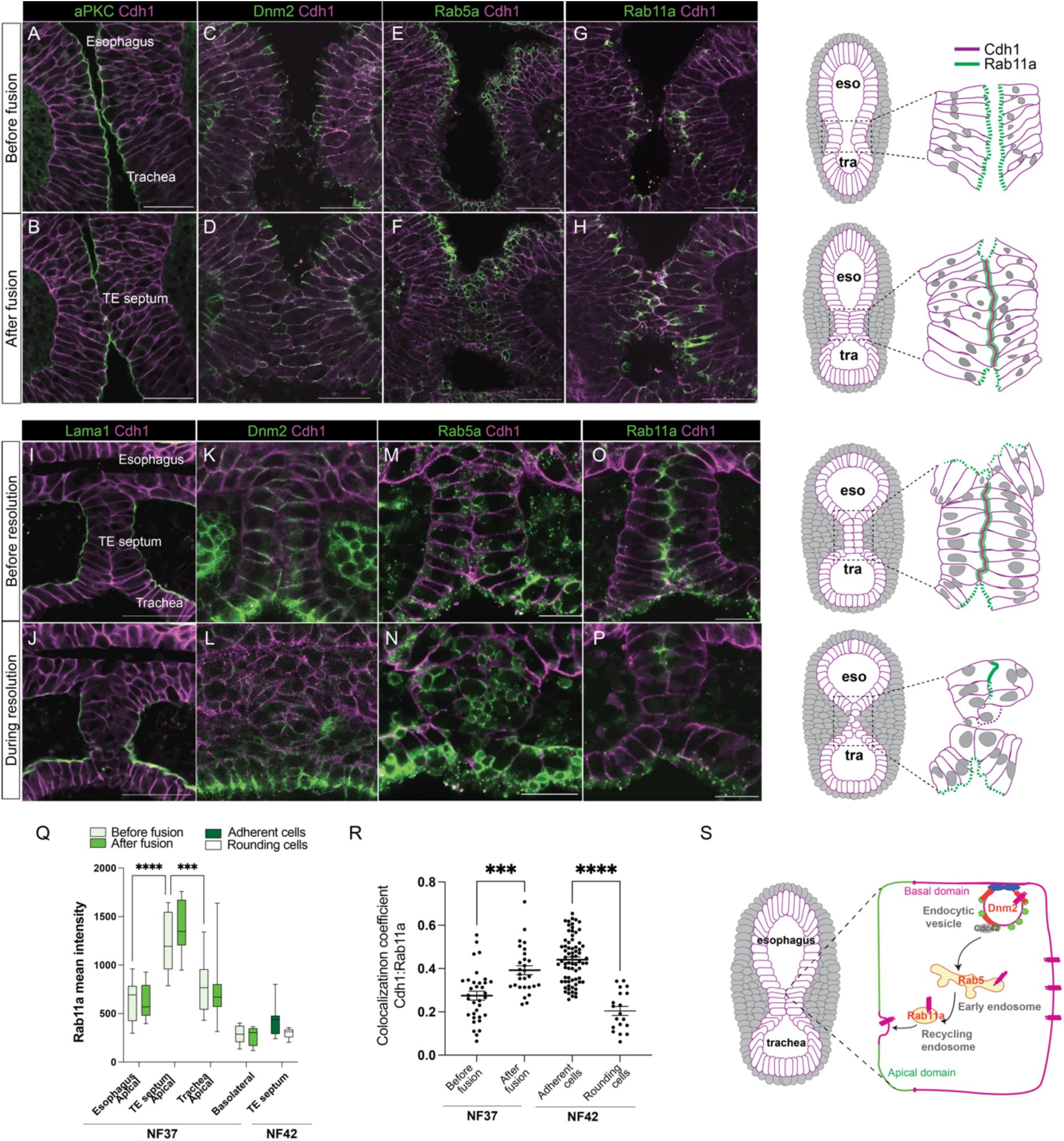
Dynamic endosome localization and epithelial remodeling during tracheoesophageal morphogenesis. A-B, I-J) Immunostaining of aPKC, Laminin (Lama1), and Cdh1 dynamics in the Xenopus foregut during trachea-esophageal morphogenesis. C-H, K-P) Immunostaining of Dnm2, Rab5a, and Rab11a in the Xenopus foregut during trachea-esophageal morphogenesis. Scale bar = 50 μm. Diagrams depict the temporal-spatial dynamics of Cdh1 and Rab11a subcellular localization during tracheoesophageal morphogenesis. Q) Quantification of Rab11a immunostaining intensity during foregut fusion and separation (mean ± min/max, 2W-ANOVA, ****p<0.0001, n=4-6 embryos analyzed). R) Cdh1-Rab11a co-localization at the epithelial interface (Mean Pearson colocalization coefficient ± SEM, *** p<0.001, **** p<0.0001 1W-ANOVA, n=5-13 cells per embryo, N=6-8 embryos per stage). S) A model of how endosome trafficking may mediate Cdh1 re-localization during epithelial fusion.

As previously reported in Nasr et al., 2019^14^, just before the foregut midline fuses at NF37, the endodermal tube is a typical pseudostratified columnar epithelium, with Cdh1 localized to the basal-lateral cell borders and in adherens junctions. Apical polarity proteins like aPKC and Par3 are localized to the luminal cell surface, whilst Laminin (Lama1) lines the basal surface, contacting the surrounding mesoderm (Figure 2A and Figure S2A-C). As the foregut constricts, apical proteins aPKC and Par3 are downregulated at the midline, where the opposing epithelia touch and Tjp1+ tight junctions are lost (Figure 2B and Figure S2D-G). This is coincident with the localization of Cdh1 to the luminal contact surface (Figure 2B, quantification in Figure 2Q), which presumably helps mediate the adhesion of the two sides and the formation of a transient, columnar epithelial bilayer by NF39 (Figure S2J). Throughout this fusion process all foregut epithelial cells (dorsal, ventral, and midline) have Dnm2 detectable at the apical-lateral membrane and subapical Rab5a+ early endosome vesicles (Figure 2C-F, quantification in Figure S2H-I). In contrast, Rab11a+ puncta are detectable at the apical membrane of epithelial cells in the foregut with higher intensity levels in the midline epithelial cells that are coming into contact (Figure G-H, quantification in 2Q).

Twelve hours later, at NF41-42, cell surface Dnm2 and Rab5a+ puncta were observed throughout the epithelial bilayer, as well as in the trachea and esophageal epithelia and the surrounding mesenchyme (Figure 2K,M). At this point, Rab11a colocalized with Cdh1 at the adhesion surface between the two epithelial sides in the bilayer (Figure 2O-R). As the transient bilayer began to resolve, we observed a downregulation of Rab11a and an increase in large Rab5a+ vesicles in the epithelium, consistent with elevated early endosome activity (Figure 2L,N,P). This coincided with a loss of Cdh1 signal on cells rounding up and losing attachment to their neighbors as the mesenchyme invaded to complete separation of the foregut (Figure S2K-L).

### Asymmetric relocalization of Cdh1 is required for trachea-esophageal separation

Our results suggested that a localized reduction in Cdh1, possibly mediated by Rab5-dependent endocytic internalization, facilitates foregut separation. To quantitatively assess Cdh1 dynamics and cell-cell contacts, we measured Cdh1 immunostaining intensity normalized to a ubiquitously expressed membrane-GFP transgene in 3D renderings of the contact surfaces between epithelial cells in the bilayer as it was separating. We found that the contact surface area between epithelial cells where the bilayer was separating was significantly smaller, with lower Cdh1 levels, compared to the larger adherent surfaces of the same cells that were still in contact with adjacent esophageal and tracheal epithelium (Figure 3A, B). This suggests that asymmetric relocalization of Cdh1 from the rounding cell surface membrane facilitates the separation of the epithelial bilayer.

**Figure 3.**
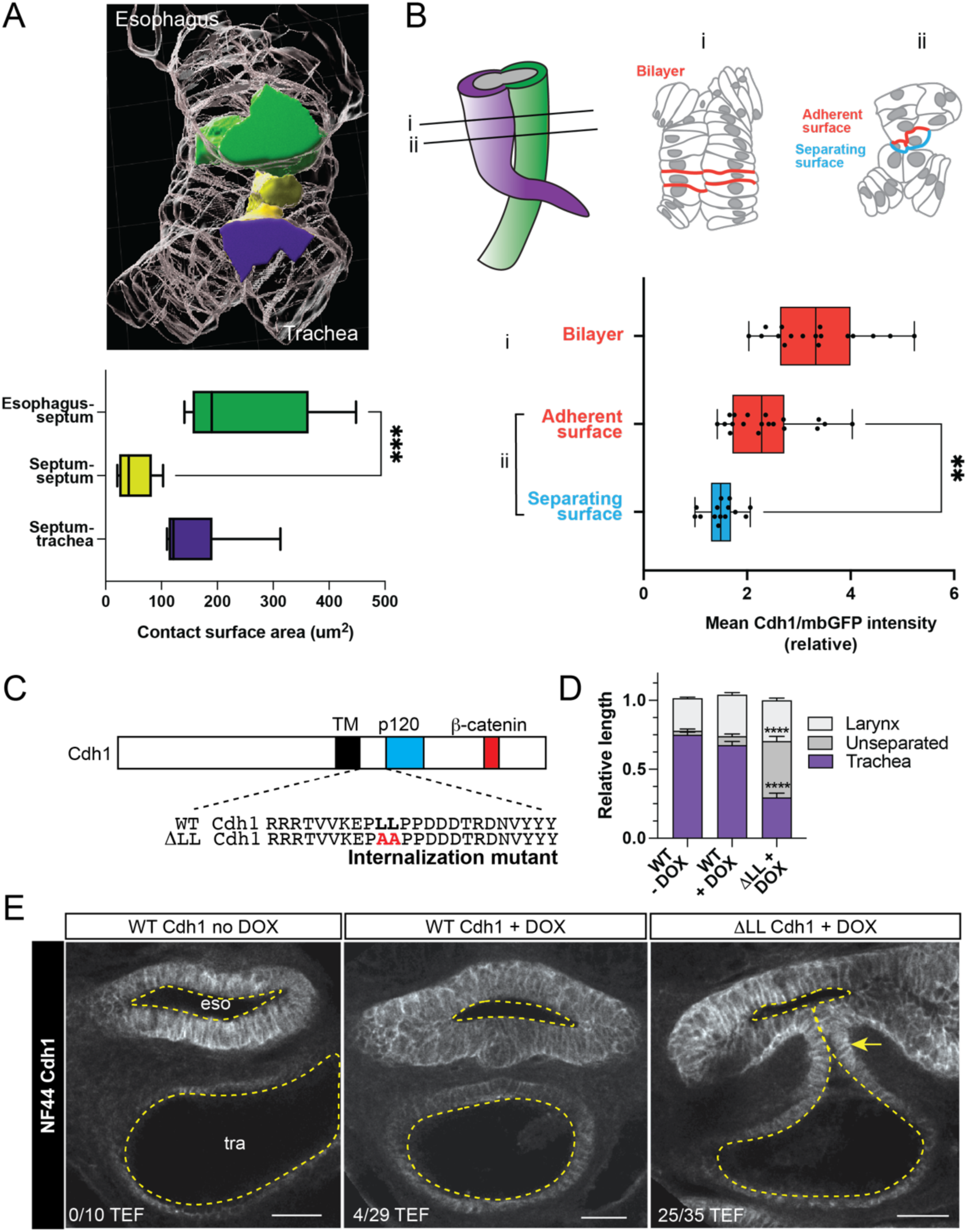
Endocytosis of Cdh1 is required for trachea-esophageal separation. A) 3D cell-surface rendering of the resolving septum; esophageal cells (green), septum cells (yellow), tracheal cells (purple). Septum cells significantly decrease surface contact area with each other compared to the contact between tracheal and esophageal cells (mean ± min/max, *** p<0.001 1W-ANOVA, n=6 cells per embryo, N=9 embryos). B) Diagrams of the separating foregut at i) the intact bilayer and ii) at the point where the bilayer is separating. The graph quantifies Cdh1/mbGFP intensity at cell surfaces in the bilayer, and the adherent versus separating side of the cells are losing contact (mean ± min/max, ** p<0.01 1W-ANOVA, n=5 cells per embryo, N=6 embryos). C) Structure of wild-type (WT) and the τιLL → AA Cdh1 mutant that cannot be internalized by endocytosis^27^. D) Quantification of the TEF phenotype in ΔLL-Cdh1 mutants compared to WT-Cdh1 and no DOX control embryos (mean ± SEM, ****p<0.0001 1W-ANOVA, n=10-35 embryos, N=3 transgenesis experiments). E) Cdh1 immunostaining and confocal microscopy of NF44 transgenic embryos and controls. Scale bars = 50 μm.

To test whether endocytic removal of Cdh1 from the cell surface was required for resolution of the bilayer, we expressed a Cdh1 mutant (ΔLL-Cdh1) that cannot be internalized by clathrin-dependent endocytosis^27^ using *Xenopus* F0 transgenics (Figure 3C). The bipartite transgene *Tg(hhex:trTA;TRE:ΔLL-Cdh1-GFP)* specifically expresses the mutant ΔLL-Cdh1 or a control wildtype (WT)-Cdh1 in the foregut upon administration of doxycycline (DOX). Compared to non-induced and WT-Cdh1 controls, approximately 70% of ΔLL-Cdh1 DOX treated transgenic embryos exhibited a TEF, with a significantly longer segment of unseparated epithelial bilayer (Fig 3D,E). This demonstrates that endocytosis of Cdh1 is required for epithelial bilayer to undergo fission and separate the trachea and esophagus.

### Endosome trafficking is required for maintenance and resolution of the epithelial bilayer during tracheoesophageal morphogenesis

Next, we determined which epithelial remodeling events driving tracheoesophageal morphogenesis require endosome trafficking. Surprisingly, initial foregut constriction and epithelial fusion, with downregulation of aPKC and increased localization of Cdh1 at the contact surface, occurred normally in *dnm2*, *rab5a*, and *rab11a* mutant embryos (Figure 4A). The earliest cellular phenotype we observed was a disorganized septum 4-6 cells wide, with cells in the middle being round rather than the stereotypical columnar epithelial bilayer at NF41 (Figure 4B, quantified in Figure 4E). Moreover, at NF42, when the septum in control embryos is separating, the mutant septum did not downregulate Cdh1 and resolve into the distinct trachea and esophageal tubes. Rather, a disorganized epithelial mass persisted, resulting in a blind-ended fistula (Figure 4C-D, Table S1). We frequently observed blisters in the mutant septa at NF42. Interestingly, the blisters exhibited asymmetric reductions in Cdh1 on the luminal surface, suggesting that some cells in the tissue try to separate from one another, but other cell-cell contacts prevent complete septation (Figure 4D). These results indicate that endosome trafficking is required to maintain an organized epithelial bilayer and suggest that the accumulation of too many cells with persistent Cdh1 at epithelial-epithelial interfaces disrupts the resolution into distinct trachea and esophageal tubes.

**Figure 4.**
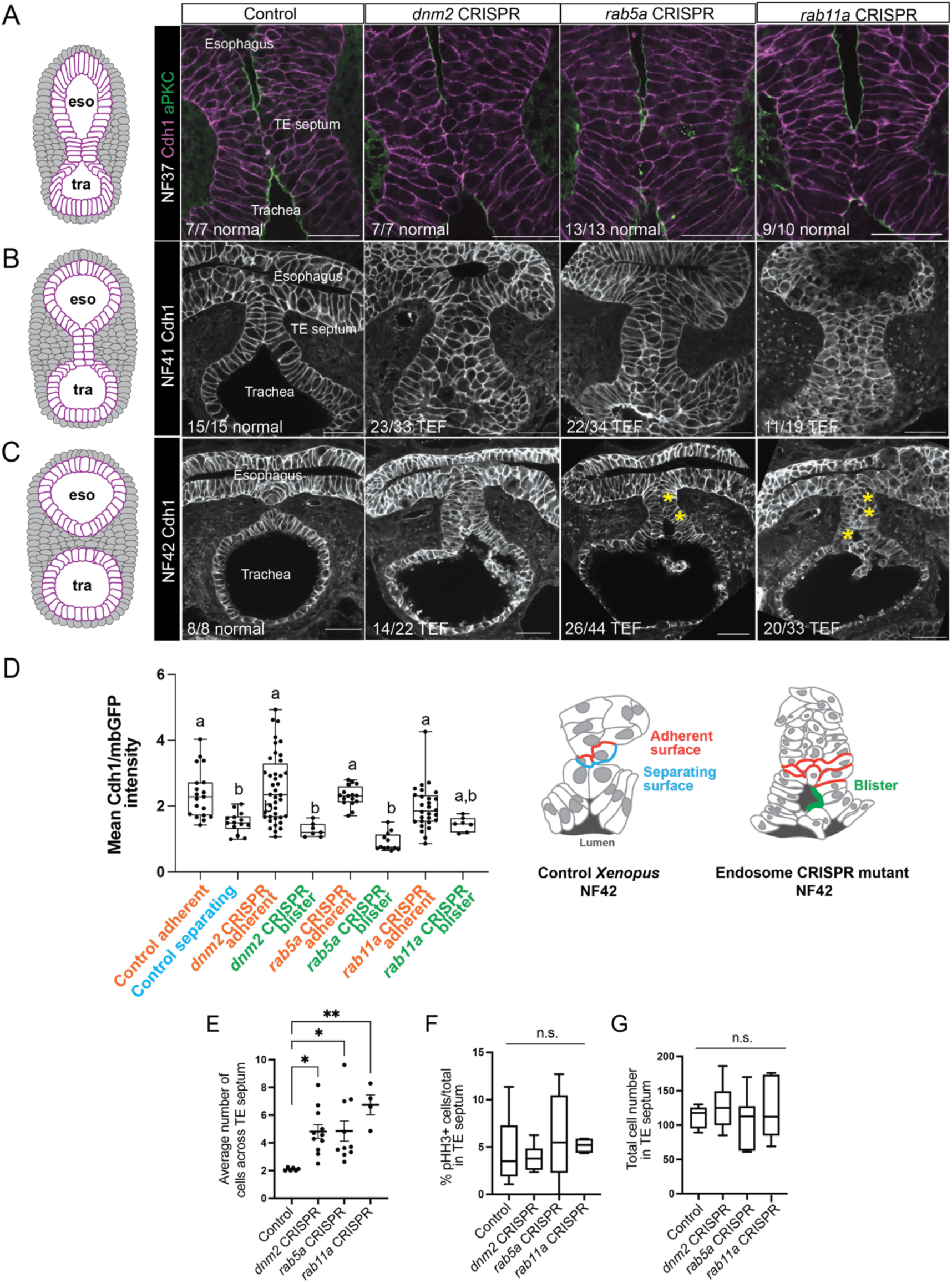
Endosome trafficking is required for resolution of the epithelial bilayer. A) At NF37, the septum in *dnm2*, *rab5a*, and *rab11a* CRISPR mutants undergoes epithelial fusion similar to controls. B) At NF41 the epithelial bilayer in endosome mutants is disorganized compared to controls. C) *dnm2*, *rab5a*, and *rab11a* CRISPR mutants exhibit TEF and blisters in the TE septum (asterisks). Scale bars are 50 µm. D) Quantification of Cdh1/mbGFP intensity in the adherent, separating, and luminal cell surfaces, in control septum, and endosome mutant blisters (mean ± min/max, 1W-ANOVA, different letters indicate p<0.05 between groups, n=7-40 cells, N=3-5 embryos) E) The NF41 mutant septa have more cells across the width compared to controls (mean ± SEM, * p<0.05 1W-ANOVA, ** p<0.01, N=6-11 embryos). F) There are no significant (n. s.) differences in proliferation (pHH3+/total) or G) total cells in the septum of mutant embryos compared to controls (mean ± min/max, 1W-ANOVA, n=46 embryos).

To determine if the accumulation of cells was due to elevated cell proliferation, we quantified confocal Z-stacks of the entire foregut region. We found that neither the total number of cells in the trachea-esophageal septum nor the rate of proliferation (as measured by phosphorylated histone H3 staining) was significantly increased in endosomal mutants compared to controls (Figure 4F-G). Three-dimensional reconstructions demonstrated that the mutant septa were wider (x-axis) and shorter in the rostral-caudal axis (z-axis) compared to controls but were similar in the dorsal-ventral axis (y-axis) (Figure S4). This indicates that mutant cells somehow accumulate across the width of the epithelia septum at the expense of fewer cells along the rostral-caudal extension of the separating trachea and esophagus.

### Endosome trafficking controls cell polarity and orientation of division in the epithelial bilayer

The earliest cellular phenotype we observed in endosome CRISPR mutants was the significant accumulation of cells inside the epithelial bilayer after foregut fusion. Since cells in pseudostratified and columnar epithelia are known to divide into the plane of the tissue to maintain appropriate organization, we considered the possibility that in mutant embryos, cells might aberrantly divide into the middle of the septum and be unable to reintegrate back into the epithelial bilayer contributing to the cell build-up (Figure 5A). To assess the plane of cell division, we measured the angle of the mitotic spindle relative to the basal membrane in the septum of endosomal mutants and controls. While cells in the epithelial bilayer of control embryos divided into the plane of the epithelia as expected, with an average spindle orientation of 0-15°, *dnm2*, *rab5a,* and *rab11a* mutants exhibited a more random distribution of mitotic spindle angles (Figure 5B,C). This defect was primarily localized to the septum, as the mitotic spindle angles in the trachea and esophagus were not significantly different between mutants and controls (Figure S4). Thus, loss of endosomal trafficking causes aberrant cell division into the center of the epithelial bilayer, and possibly a defect in epithelial reintegration into the tissue after cell division due to altered apical-basal cell polarity.

**Figure 5.**
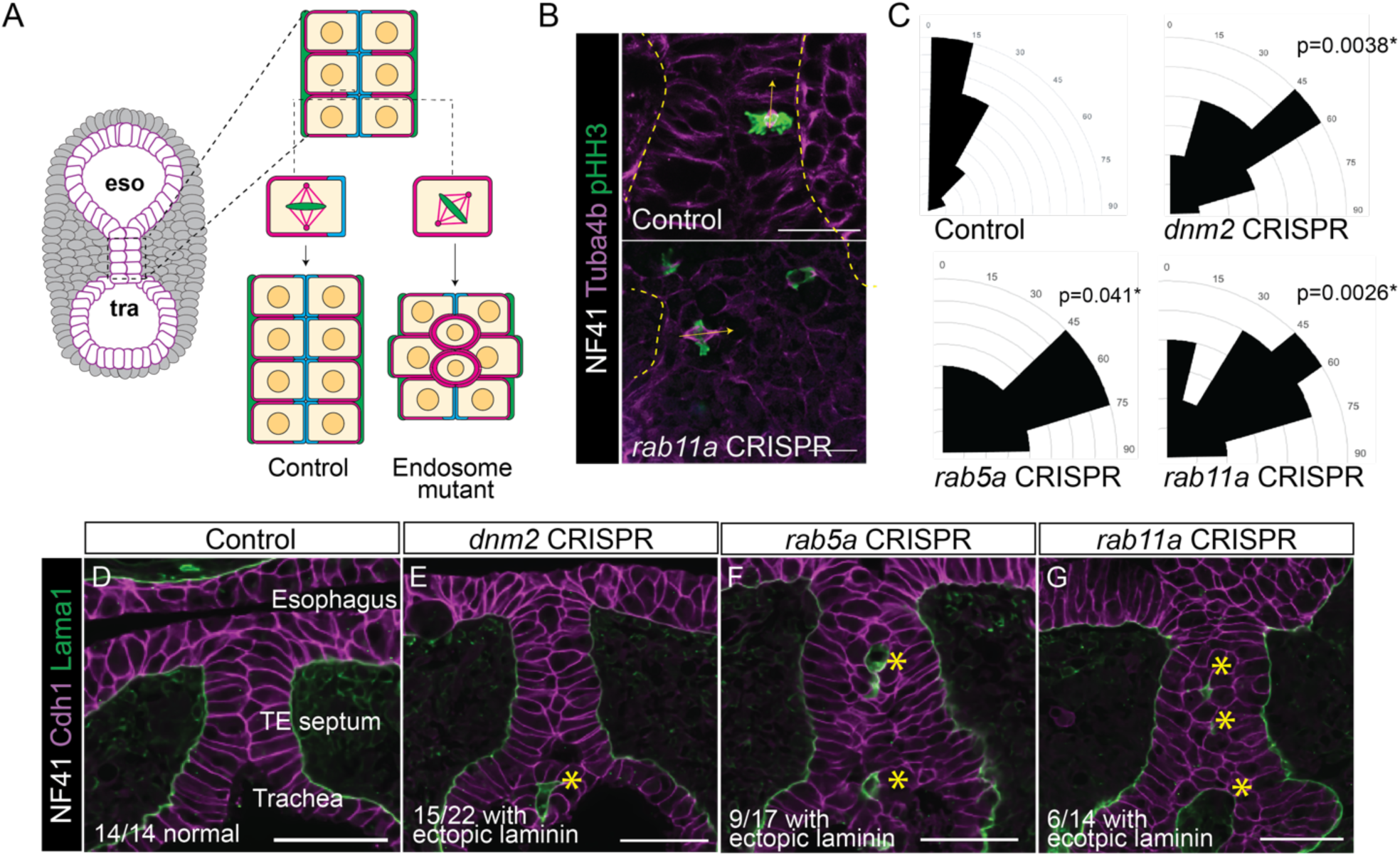
Endosome trafficking controls the polarity of epithelial cell division. A) A model of altered cell division orientation in endosome mutants causing cells to accumulate in the epithelial septum. B) Immunostaining of the mitotic spindle (Tuba4a and phospho-histone H3) was using to measure the angle of cell division at NF41. C) Distribution of mitotic spindle angles is random in endosome mutant epithelial cells compared to controls, which consistently divide between 0-15° (n=3-15 spindles per embryo, Kolmogorov-Smirnov test, N=4-6 embryos. D-G) Endosome mutants have ectopic Laminin deposits (asterisks) inside the blistered disorganized epithelia. Scale bars are 50 µm.

To test whether apical-basal cell polarity was disrupted, we performed immunostaining for laminin, which is secreted from the basal side of epithelia into the extracellular matrix. Indeed, mutant embryos exhibited ectopic laminin deposition in the center of the disorganized septum surrounding the round central cells and in the lumens of blisters within the tissue (Figure 5D-G). These observations are consistent with disrupted endosome trafficking leading to loss of apical-basal polarity in the transient epithelial bilayer, randomized cell division, and cells accumulating in the mutant tracheoesophageal septum.

### Endosome trafficking localizes Vangl2 to maintain apical memory in the epithelial bilayer

During foregut morphogenesis, when the apical epithelial cell surfaces from either side touch one another, they lose apical markers (aPKC, Par3) and gain Cdh1 (typically basal-lateral) as they adhere and form the transient bilayer. Despite this apparent loss of apical identity, the cells must somehow retain some apical-basal cues to maintain an organized bilayer and undergo cell division in the plane of the epithelium – a process that we found requires endosome trafficking. One candidate pathway that might regulate this process is the Vangl-Celsr polarity complex. In addition to their well-known role in the Wnt/PCP pathway where heterotypic cell-cell interactions between transmembrane Vangl, Celsr, and Frizzled proteins govern cell orientation in the plane of the epithelia^28^, Vangl and Celsr are also involved in specifying apical-basal polarity in epithelia by recruiting apical Par3 and aPKC whilst restricting Scrib to the basolateral domain^18,29,30^. Furthermore, apical Vangl-Celsr complexes are reported to be internalized by endosomes during polarized cell divisions of the fetal mouse epidermis^31^, suggesting a potential mechanistic link in the foregut.

To assess whether Vangl-Celsr complexes are involved in endosome-dependent epithelial remodeling in the foregut, we performed Vangl2 and Celsr1 immunostaining in combination with Rab11a (antibodies are not available for *Xenopus* Celsr2 and Vangl1). Before foregut fusion, diffuse Vangl2 was observed throughout the foregut epithelia with no apparent difference between apical and basolateral cell domains (Figure 6A,F). However, as the midline epithelia touched, Vangl2 became progressively co-localized with Rab11a puncta at the luminal/apical contact point and subsequent adherent interface of the bilayer (Figure 6, Figure S6). Super-resolution microscopy demonstrated that approximately 40% of Rab11a and Vangl2 puncta at the adherent interface of the bilayer were within 500 nM of each other, consistent with the size of a recycling endosome vesicle^32,33^. In contrast to this dynamic Vangl2 subcellular localization, Celsr1 remained around the entire juxta-membrane region of the epithelial bilayer cells throughout tracheoesophageal morphogenesis (Figure S6E-H).

**Figure 6.**
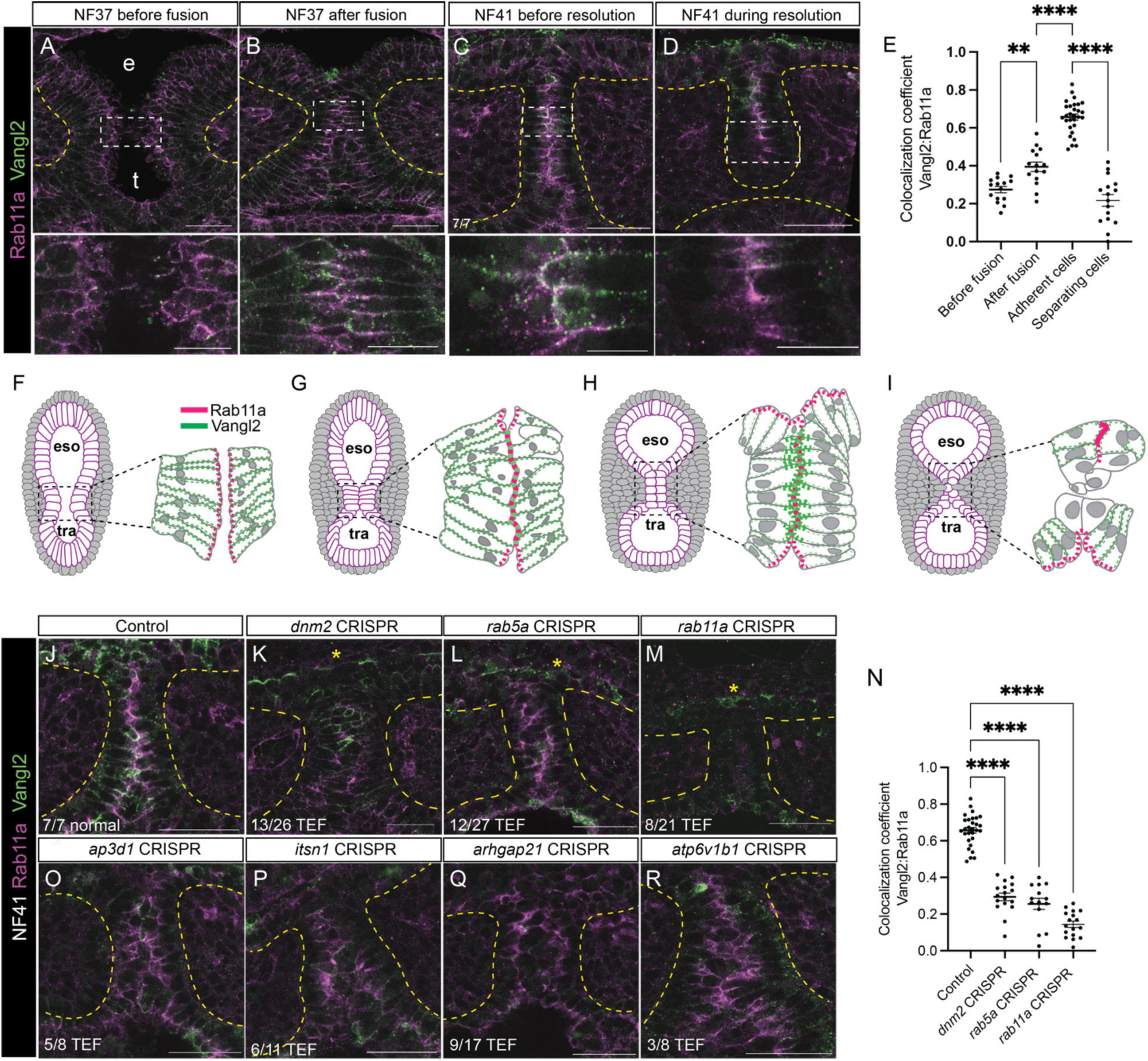
Recycling endosomes localize Vangl2 to maintain apical memory in the transient trachea-esophageal septum. A-D) Time course of Vangl2-Rab11a immunostaining during *Xenopus* tracheoesophageal separation. E) Dynamic Vangl2-Rab11a co-localization at the epithelial interface (Mean Pearson colocalization coefficient ± SEM, *** p<0.001, **** p<0.0001 1W-ANOVA, n=6 cells per embryo, N=5 embryos per stage). F-I) Diagrams of Vangl2-Rab11a localization during foregut separation. J-M) Vangl2 is mis-localized in septum epithelial cells of endosome mutants. N) Endosome mutant septa have significantly lower Vangl2-Rab11a colocalization relative to controls (mean ± SEM, **** p <0.0001 1W-ANOVA, n=5 cells per embryo, N=4-6 embryos per condition). O-R) CRISPR F0 mutation of patient variant orthologous genes in *Xenopus* (*ap3d1, itsn1, arhgap21*, and *atp6v1b1)* disrupts Vangl2 localization in the TE septum. Scale bars are 50 µm (20µm in insets).

Considering that endosomes are reported to internalize cell surface Vangl2 in the fetal mouse epidermis^26^, this suggested that Vangl2 might be a cargo of the Rab11+ recycling endosomes, transporting it to the cell surface where the epithelia fuse. To test this possibility, we examined Vangl2 localization in the endosome mutants. Vangl2-Rab11a co-localization at the epithelia contact was lost in the disorganized multilayer septum of *dnm2*, *rab5a* and *rab11a* mutants (Figure 6J-N, Figure S6D). However, Celsr1 localization and apical Vangl2 in the esophagus and trachea were unaffected (Figure 6K-M, asterisks, Figure S6I-M), indicating that recycling endosomes are required to localize Vangl2 at the midline apical interface.

### Mutation of EA/TEF patient-like genes in *Xenopus* disrupts Rab11-Vangl2 localization

We next assessed whether a similar mechanism was involved in the TEF observed when we mutated the *Xenopus* orthologs of endosome pathway risk genes from patients. We tested *AP3D1*, *ARHGAP21*, and *ITSN1* which encode adaptors and CDC42-GEFs respectively during endocytic vesicle formation, as well as *ATP6V1B1*, which encodes a vacuolar ATPase subunit required for acidification of endosomal vesicles^34–38^. As predicted, F0 CRISPR mutants of *ap3d1*, *arhgap21*, *atp6v1b1* and *itsn1* all exhibited a loss of apical Vangl2 localization and a disorganized persistent septum (Figure 6O-R), consistent with these proteins functioning in the recycling endosome pathway.

Together these results indicate that during tracheoesophageal morphogenesis, dynamic recycling endosome activity is required to localize Vangl2 at the cell surface where the opposing midline epithelia adhere to one another. Vangl2 appears to define an “apical memory” in the epithelial bilayer cells which have lost aPKC and upregulate Cadherin, a membrane configuration more typical of adherent basal-lateral cell-cell contacts. This unexpected behavior is distinct from the typical roles for Vangl2 in apical-basal and planar cell polarity.

### Loss of Vangl or Celsr disrupts tracheoesophageal morphogenesis resulting in TEF

The results above imply that Vangl-Celsr is required for tracheoesophageal morphogenesis. Interestingly, one patient from our EA/TEF cohort had a heterozygous *de novo* damaging missense variant in *CELSR2* predicted to alter an amino acid (N953S) in an N-terminal cadherin domain^23,39^. Moreover, a rare *de novo* variant in *CELSR1* (V1452I) was previously reported in another EA/TEF patient^40^. Together with our observations that cell polarity is disrupted in endosome mutants, this led us to test whether Vangl-Celsr were required for trachea-esophageal morphogenesis in *Xenopus*.

Loss of function CRISPR-Cas9 F0 *Xenopus* mutants for *celsr2*, *celsr1,* or *vangl2* all resulted in variably penetrant TEF in 25-30% of the embryos at NF44 (Table 1). Examination of the mutant foreguts at NF40 revealed a disorganized, thicker septum with round cells in the middle and frequent blisters, phenocopying the endosome mutants (Figure 7A-D). Moreover, the *celsr1 and celsr2* mutants exhibited reduced Vangl2 immunostaining, consistent with previous reports that Celsr1 forms a heterotypic cell-cell complex with Vangl2^28^. Interestingly, in *vangl2* and *celsr1/2* mutants Rab11a was still localized to the middle of the bilayer (Figure 7E-H), indicating that Vangl-Celsr complexes are required for tracheoesophageal morphogenesis downstream of recycling endosomes. Together, these results suggest that an unexpected but evolutionarily conserved endosome-cell polarity pathway controls tracheoesophageal morphogenesis and that disruptions to this process might contribute to EA/TEF in many patients (Figure 7I).

**Fig. 7.**
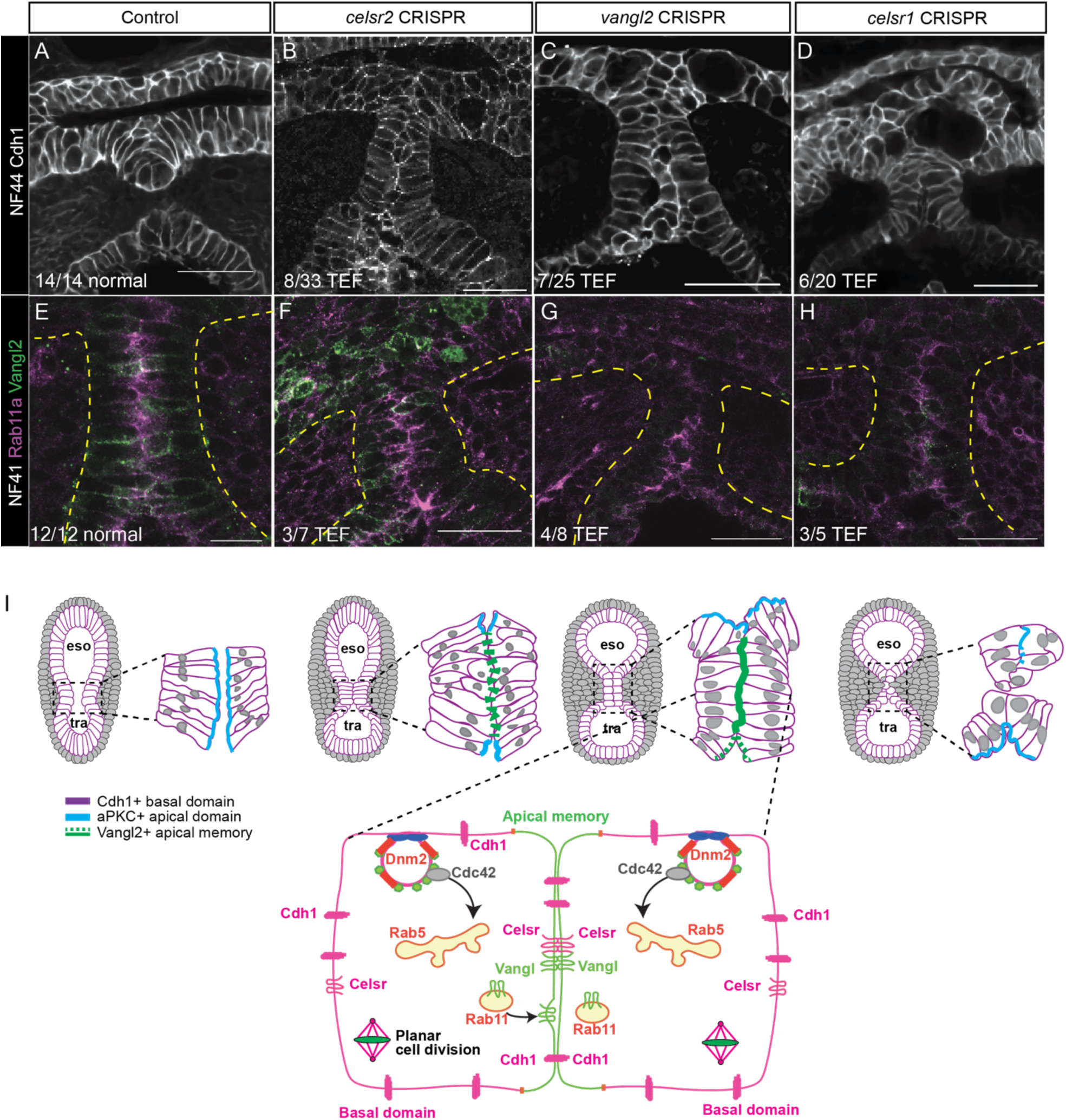
Loss of Vangl or Celsr disrupts tracheoesophageal morphogenesis, resulting in TEF. (A-D) Cdh1 immunostaining in *celsr2*, *vangl2*, and *celsr1* mutants with TEF (E-H) disorganized epithelial septa that fail to co-localize Vangl2-Rab11a. Scale bars are 50 µm. I) Model of how endosome trafficking of Vangl2 maintains apical-basal polarity information and the proper tissue organization required for tracheoesophageal morphogenesis.

## Discussion

In this study, we demonstrate that an endosome-cell polarity pathway is required for the epithelial remodeling that drives *Xenopus* tracheoesophageal morphogenesis. We discovered that Rab11+ recycling endosomes localize the transmembrane polarity protein Vangl2 to the contact surface where foregut epithelia fuse. This appears to confer an “apical memory” which is essential to maintain an organized transient epithelial bilayer and its subsequent fission into distinct trachea and esophagus (Figure 7I). This unexpected Rab11-Vangl2 activity is distinct from its typical roles in planar cell polarity and apical basal polarity, and may represent a general paradigm for epithelial fusion and fission during morphogenesis.

Our study provides new insight into how cell and tissue polarity is regulated during tissue fusion and fission through endosome-mediated dynamic relocation of polarity and cell adhesion proteins. Previous work suggests that the hierarchy of endosome trafficking and cell polarity in development is context dependent. In the fetal mouse epidermis, endosomes internalize Vangl, Celsr, and Fzd proteins organized along the proximal-distal axis in typical PCP fashion, which are later relocated back to the appropriate cell surface after cell division to maintain epithelial organization^31^. In contrast, Vangl2 functions upstream of Rab11 in classical Wnt/PCP and apical-basal polarity roles to regulate convergent extension, planar cell polarization and apical constriction during Xenopus gastrulation and neural tube morphogenesis^41–43^. We show in the foregut that Rab11a is upstream of Vangl2, as Rab11a is still localized properly in vangl/celsr F0 CRISPR mutants. We propose that Vangl2 confers “apical memory” to cells that have otherwise lost apical character. The Vangl-Vangl interface is unexpected since in most other reports, Vangl-Celsr interactions are heterotypic, with Vangl+ cells binding to Celsr+ / Vangl-cells. In addition, we find no evidence of Vangl-mediated PCP activity during tracheoesophageal morphogenesis. Another distinction between what we have described in the foregut and these other examples of Vangl-regulated epithelial morphogenesis is that in the foregut the epithelia first fuses (like the neural tube and palate) but then undergo fission. Other tissues that experience this type of dynamic fusion-fission morphogenesis include separation of anorectal tubes in the hindgut and semicircular canals in the ear^44–46^. We predict that similar Rab11a- Vangl2 mechanisms may act in those tissues.

We also show a role for Vangl2 that is distinct from its typical functions in apical-basal polarity. Our observation that Vangl2 and Rab11a colocalize coincident with a loss of Par3 and aPKC in the foregut bilayer epithelium was unexpected since Vangl2 is typically thought to help recruit these proteins to the apical surface in a polarized epithelium. There is precedence for Vangl2 activity independent of its classical polarity roles. In the developing mouse pancreas, apical restriction of Vangl2 maintains epithelial integrity independent of apical-basal cell polarity^47^. Vangl2 also has cell shape and polarity functions in the developing mouse lung mesenchyme during branching morphogenesis independent of its PCP or apical-basal polarity roles^48,49^. Neither the *looptail* mutation (dominant-negative) nor conditional knockout of Vangl2 using ShhCre impairs trachea-esophageal separation. We postulate that a different Cre driver with earlier recombination in the foregut might lead to separation anomalies. The observation that endosomal mutants have a TEF penetrance of ∼60% while Vangl-Celsr F0 mutants only have a 30% penetrance suggests there may be genetic redundancy (Rab11a vs Rab11b) and Vangl family members (Vangl1 vs Vangl2). Redundancy between Rab11a/b was shown in the developing mouse pancreas, as knockout of both Rab11a and Rab11b was required to see epithelial defects^50^. Another possibility is that in addition to Vangl2, other endosome cargo proteins participate in foregut morphogenesis.

Cadherin is a candidate cargo of recycling endosomes in the foregut. Unlike Vangl2, Cdh1 can still localize to the midline of the fused epithelial bilayer in endosome mutants. However, we show that Cdh1 needs to be internalized by endocytosis at the separating epithelial cell surface for fission of the bilayer into the trachea and esophagus. We postulate that other endosomal cargo may include Integrins that facilitate tissue adhesion and Laminins that mediate interaction with the adjacent mesoderm at the basal surface. Consistent with this possibility, we note that EA/TEF patients were identified with potentially damaging variants in *ITGB4* and *LAMB4*^23^. Another candidate pathway that might regulate endosome-mediated loss of adhesion is the Eph/Ephrin pathway. Eph/Ephrin are known to regulate tissue separation and cell sorting in many contexts^51^ and *Efnb2* was recently shown to be required for trachea-esophageal separation downstream of Nkx2-1^52,53^. In the future we will test the intriguing possibility that Ephrin-Cadherin-endosome-Vangl interactions mediate tissue adhesion and separation in the developing foregut.

Endosome trafficking is a ubiquitous cell pathway required for homeostasis and many basic cellular functions. Several pleiotropic diseases are associated with variants in endosome trafficking genes^22^, but interestingly many of these are rather specific to certain tissues. Not surprisingly, total loss of endosome activity leads to early embryonic lethality in germline mutant mice^54,55^ and in *Xenopus* embryos results in gastrulation and neurulation defects^41^. Our F0 CRISPR mutagenesis on the other hand results in 58-90% indel mutations, more damaged than a heterozygote but not complete nulls. Indeed, in our experiments, endosome mutant embryos have a significantly higher frequency of gastrulation and neurulation defects compared to controls where the pigmentation gene *tyr* was mutated. In our experiments we selected those embryos that survived to later stages. The fact that later embryos with partial loss of endosome activity show specific defects suggests that developing organs and tissue undergoing dynamic epithelial remodeling, like tracheoesophageal separation, represent developmental bottlenecks that are particularly sensitive to disruptions in endosomal trafficking. This may partially explain why EA/TEF patients frequently present with variants predicted to have partial loss of function or hypomorphic variants in endosome and polarity pathway genes.

Our results are consistent with the “saddle rising” model of trachea-esophageal separation proposed by Que and colleagues^4^ and provide a cellular basis for the saddle as the point of epithelial fusion-fission and mesenchymal invasion. We note differences between *Xenopus* and mouse TE morphology and anatomy – in mice, the epithelial fusion event is more transient and only 3-4 cells in length, while in *Xenopus*, the epithelial fusion event is more prolonged, forming a bilayer septum tissue over 10 cells in length^14^. In mammals the foregut tissue significantly grows in the longitudinal plane as the trachea and esophagus separate resulting in an apparent “rising” of the saddle compared to *Xenopus*. Our study uses *Xenopus* as a model to perform rapid disease causality screening and cellular analysis to minimize the confounding issue of tissue growth.

The EA/TEF risk genes validated in this study are involved in various stages of endosome trafficking, including vesicle formation (ITSN1, AP1G2), cytoskeletal transport (ARHGAP21, ABRA, RASAL2, RAPGEF3), vesicle acidification (ATP6V1B1) ^23,40^. Interestingly the patients enriched in endosome variants frequently had co-morbidities similar to those observed in patients with VANGL1/2 mutations ^28^ such as neural tube anomalies^56,57^, renal anomalies^58,59^, congenital heart anomalies^60,61^, and lung diseases^62^. While further studies are required to test a causal link between endosome and polarity proteins in patients, it is tempting to speculate that complex EA/TEF patients tend to have comorbidities in organs that experience tissue fusion-fission during fetal development^23^. This suggests the provocative possibility of an “endosomeopathy” syndrome, similar to ciliopathies. Further comparative analysis of patient genetics and animal models will deepen our understanding of tissue fusion and fission events during development, aiding the care and treatment of patients.

### Limitations of the study

The F0 CRISPR-Cas9 gene editing strategy used in our study is inherently variable compared to analyzing germline mutants. We took advantage of rapid *Xenopus* embryonic development to efficiently screen patient risk genes generating simple indels that produced loss of function mutations. However, it is possible that variants in EA/TEF patients result in missense gain of function or hypomorphic proteins. Moreover, the available evidence suggests that EA/TEF risk is polygenic and our approach does not account for this. Indeed, we found that some of the gene mutations resulted in a low penetrance TEF, which was not significant over controls, but might indicate a contribute to the phenotyope in a polygenic state. Future studies should model the patient’s variants and consider multiple genes acting together.

We also do not rule out the possibility that endosome trafficking is required in the surrounding mesenchyme. This might include polarized secretion of matrix metalloproteases for basement membrane degradation or PCP-mediated directional mesenchyme migration across the midline during trachea-esophageal separation. Follow up studies using tissue-specific manipulations will be necessary to address this possibility.

## Acknowledgements

We thank the Bio-Imaging and Analysis Facility for imaging and analysis advice (RRID:SCR_022628), the Division of Veterinary Services, and the DNA Sequencing Core at CCHMC (RRID:SCR_022630) for their services used in this study. We thank S. Sokol for kindly providing the anti-Vangl2 antibody. We thank the resources provided by the National Xenopus Resource (RRID:NXR_0012) and Xenbase (RRID:SCR_003280)^63^. We thank the members of the CLEARconsortium.org for access to EA/TEF patient data and for helpful discussions. This work is supported by NICHD P01HD093393 to YS, WKC, and AMZ, and a Canadian Institutes of Health Research postdoctoral fellowship to NAE. This project was supported in part by NIH P30 DK078392 of the Digestive Diseases Research Core Center in Cincinnati for the Bio-Imaging and Analysis Facility and the DNA Sequencing Core at CCHMC.

## Author contributions

Conceptualization: NAE, AMZ. Formal analysis: NAE, AK, AMZ. Funding acquisition: YS, WKC, AMZ. Investigation: NAE, SAR, AK, AW, ZA, APK. Methodology: NAE, SAR, MJK. Project administration: NAE, AMZ. Supervision: NAE, AMZ. Validation: NAE. Visualization: NAE, AMZ. Writing – original draft: NAE, AMZ. Writing – review & editing: all authors.

## Methods

### Experimental models

*Xenopus* experiments were performed using Cincinnati Children’s Hospital Medical Center (CCHMC) Institutional Animal Care and Use Committee (IACUC 2022-0026) approved guidelines. Wild type adult *Xenopus tropicalis* and *Xenopus laevis* frogs were purchased from NASCO and Xenopus1 (USA) and housed in the CCHMC vivarium under standard housing conditions. *Xenopus* embryos were obtained by *in vitro* fertilization according to standard procedures^64^. Transgenic membrane GFP *Xenopus laevis* (*Xla.Tg(CMV:memGFP;cryga:mCherry)^NXR^*) were purchased from the National Xenopus Resource (RRID:NXR_0012). *Sox2* germline mutant embryos were generated at the National *Xenopus* Resource according to standard procedures (RRID:SCR_013731).

### CRISPR-Cas9 gene editing and Xenopus embryo culture

Guide RNAs and genotyping primers were designed using CRISPRScan, InDelphi, and CRISPOR predictive algorithms^65^ and synthesized as altR-cRNA by IDT (sequences in Resources Table and Table S3). Guide RNA sequences that maximized predictive efficiency (CRISPRScan) and high percentage of predicted out of frame indel mutations (InDelphi) were selected. After annealing with tracrRNA to create a single gRNA, gRNAs were combined with recombinant Cas9 protein (PNA Biosciences) and injected into 1-2 cell stage *Xenopus* embryos at a final concentration of 500-700pg gRNA and 1-1.5ng Cas9p. *Xenopus tropicalis* embryos were raised at 25°C in 0.1X Modified Barth’s solution (MBS) for three days post fertilization before being anesthetized and fixed at NF44 in MEMFA (3.7% formaldehyde, 0.1M MOPS, 2mM EGTA, 1mM MgSO_4_) for 1hr at room temperature, then in Dent’s solution (80% methanol, 20% DMSO) for long term storage at −20°C. *Xenopus laevis* embryos were raised between 15-21°C in 0.1X MBS for 3-5 days post fertilization (to stages NF37-44), then anesthetized and fixed in MEMFA or 2% trichloroacetic acid (for endosome proteins and Vangl2 immunostaining) for 30- 40 mins at room temperature. Embryos were immunostained and imaged by confocal microscopy as outlined below prior to genotyping.

### Genotype-phenotype scoring

Genotyping for CRISPR-Cas9 editing efficiency in individual CRISPR-edited embryos was performed as described in Zhong et al., 2022^23^ using the primer sequences in Resources Table and Table S3. PCR amplified products from individual embryos were sent for Sanger sequencing at the DNA Sequencing Facility at CCHMC. Sanger sequences were deconvoluted using Inference of CRISPR Edits (Synthego, USA) or Tracking of Indels by Decomposition, TIDE, Brinkman et al., 2014^66^) to determine percent indel efficiency (percent of mutations in each embryo) and mosaicism (different types of indel mutations in a given embryo) (Figure S1). Only embryos with >40% indel efficiency were considered in genotype-phenotype analysis, as if the mutation burden was less than this, we concluded that the gene was not sufficiently mutated to test its function. A minimum of 20 individual mutant embryos from 3 independent editing experiments were analyzed to identify a statistically significant frequency of trachea-esophageal anomalies by Student’s t-test from the 2% baseline detected in controls. Individual genotype-phenotype scoring and mutation efficiency are provided in Table S2.

### E-Cadherin site-directed mutagenesis and *Xenopus* transgenesis

pCS2+ *Xenopus* E-cadherin-3xGFP was a gift from Ann Miller (University of Michigan) and described in Higashi et al 2016^67^. A dileucine motif (LL) in the evolutionarily conserved juxta-membrane cytoplasmic domain of E-cadherin is required for E-cadherin endocytosis^27^. Site-directed mutagenesis was performed with the GeneArt Mutagenesis kit (ThermoFisher A13282 with Accuprime Taq Polymerase, ThermoFisher12339016) using the following primers (mutant sequence bold, underlined; wt sequence TTA CTA encodes LL, mutation GCA GCG encodes AA to mutate these two leucines (LL) at positions 732, 733 into alanines (AA) of Xenopus E-cadherin.

Forward mutagenic primer:

5’- GTG GTA AAA GAG CCT **<u>GCA GCG</u>** CCT CCA GAA GGC GAC −3’

Reverse Mutagenic primer:

3’- CAC CAT TTT CTC GGA **<u>CGT CGC</u>** GGA GGT CTT CCG CTG −5’

Mutagenesis was confirmed with the following sequencing primers:

Forward sequencing primer: 5’ GAG CTG GAG ATT GGT CAA TAC G 3’;
Reverse sequencing primer: 5’ GAG CCA CTG CCT TCA TAA TCG 3’.

Following confirmation of mutagenesis, both the Cdh1-GFP wild type and mutant (LL>AA) were cloned into the pCR8-Gateway TOPO vector (ThermoFisher K250020) according to manufacturer’s instructions. Gateway LR Clonase II Plus (Thermo Scientific #12538120) was then used in standard recombination reactions according to manufacturer’s instructions to transfer the *EcadGFP wild type and mutant (LL>AA)* into the pDXTRE transgenesis responder plasmid^68^, which enables Dox-dependent expression.

Transgenic embryos were generated as follows: 250pg of pDXTP-*hhex* promoter:rtTA (Nasr et al., 2019^14^) and 500pg of pDXTR-EcadGFP (wt or LL>AA mutant) were incubated in a 25uL reaction containing 2.5μL of I-SceI meganuclease enzyme (New England Biolabs #R0694S) in 0.5X I-SceI buffer. The reaction was incubated at 37^°^C for 40 minutes and then 5nL of this reaction mix was immediately injected into 1-cell embryos on either side of the sperm entry point within the first 60-75 minutes after fertilization. Embryos were cultured at 13-15^°^C for the first two to three hours after injection and subsequently at 18^°^C to 23^°^C degrees thereafter. Transgenic embryos were selected based on GFP fluorescence in the eye, which becomes visible during early tailbud stages^68^. Transgenes were activated by the addition of doxycycline (Sigma #D9891) at a final concentration of 50μg/mL to the embryo culture buffer (0.1X MBS). Culturing of un-injected embryos in this dose of Dox has no noticeable effect on embryonic development nor tracheal-esophageal separation. Transgenic efficiency (embryos with GFP+ eyes at tailbud stages NF33-35) ranged from approximately 8% to 16%, consistent with previous transgenic studies^14,68^.

### Whole mount immunostaining and confocal microscopy

Whole mount immunostaining on *Xenopus* tissue was performed as described in Nasr et al., 2019^14^ and Zhong et al., 2022^23^ with the following modifications for imaging endosome and polarity proteins: fixed embryos were rehydrated in PBS, embedded in 2% low melt agarose in PBS, and sectioned with a Vibratome VT1000S (Leica) to obtain 200 µm tissue slices containing the trachea and esophagus. Slices were permeabilized with 0.1% Triton X-100 in PBS (0.3% Triton X-100 for anti-Vangl2), blocked in 5% normal donkey serum in PBS, then incubated in primary antibody (1:100) in blocking solution overnight at 4°C. Slices were incubated in Alexa fluor-conjugated secondary antibody (1:500) in blocking solution for 2 hours at room temperature, then dehydrated in 100% methanol. For confocal microscopy, slices were cleared in BABB and imaged on a Nikon A1 inverted laser scanning confocal microscope with a 10X, 20X water immersion, or 40X oil immersion objective (Nikon USA). For super resolution microscopy, slices were cleared in BABB and imaged on a Nikon AXR confocal microscope with spatial array detector (NSPARC, Nikon, USA) in super resolution mode with a 40X and 100X oil immersion objective.

### Image analyses

#### Signal intensity quantification

Signal intensity quantification was performed in ImageJ (NIH). Regions of interest (ROI) were drawn around the apical and basal membranes, then mean gray value was measured and plotted as mean intensity ± max/min normalized to the fluorescence intensity of memGFP signal. At least five cells were measured per embryo in a minimum of five embryos per condition for quantification and statistical analysis by 2W-ANOVA.

#### Prolate ellipticity, surface contact area, and 3D quantification of Cdh1

Prolate ellipticity and surface contact area measurements were performed in Imaris (Bitplane). 3D cell volumes were rendered based on E-Cadherin staining and surface models were generated. Prolate ellipticity was measured from rendered surfaces of epithelial cells and plotted as mean ± max/min. Values approaching 1 have elongated shapes while values approaching 0 have spherical shapes. Surface-surface contact area between the esophagus and trachea-esophageal septum, within the trachea-esophageal septum, and between the trachea and trachea-esophageal septum was measured from rendered surfaces of epithelial cells using the Surface-Surface Contact Area XTension and plotted as the mean ± max/min. Mean Cdh1 intensity was measured on the rendered surface. At least five cells were measured per embryo in a minimum of five embryos per condition for quantification and statistical analysis by unpaired Student’s t-test (prolate ellipticity) and 1W-ANOVA (surface contact area).

#### Three-dimensional measurements of axes in the foregut

Three dimensional lengths of x (medial), y (dorsal-ventral), and z (rostral-caudal) axes were performed in Imaris (Bitplane). 3D cell volumes were rendered based on E-Cadherin staining and surface models were generated. Lengths were measured from rendered surfaces of epithelial cells and plotted as the mean length of each axis ± standard error of the mean. At least five embryos were measured for quantification and statistical analysis by unpaired Student’s t-test.

#### Mitotic spindle orientation

Mitotic spindle orientation angles were obtained in ImageJ (NIH) as described in Franco and Carmena, 2019^69^. Only images with z-sections through the axial axis of *Xenopus* embryos containing clearly identifiable mitotic spindles stained with anti-⍺-tubulin were analyzed for spindle angles. One line of the angle was drawn through linking both centrosomes, and the second line of the angle was drawn parallel to the basement membrane of the epithelial cell. Measurements were collected and wind rose plots were generated in Excel. At least three mitotic spindles were measured in a minimum of 5 embryos for quantification and statistical analysis was performed by Kolmogorov-Smirnov tests.

#### Colocalization and nearest neighbor analysis (super resolution)

Pearson colocalization coefficients between Vangl2/Rab11a, Cdh1/Rab11a, and Celsr1/Rab11a immunostaining signals were calculated in Nikon Elements (Nikon USA). ROIs were drawn around an entire cell, and the Pearson colocalization coefficient was determined. A coefficient of 1 indicates two channels whose fluorescence intensities are perfectly linearly related. For super resolution imaging quantification, Rab11a and Vangl2 spots were detected and rendered in Imaris (Bitplane USA). Distances between Rab11a-Vangl2 spots were measured and graphed as a histogram. Vangl2 spots within 500nm of Rab11a spots were considered to be within the Rab11a+ recycling endosome compartment^32,33^. At least six cells were measured in a minimum of 5 embryos for quantification and statistical analyses by 1W-ANOVA.

### Immunoblotting

Whole Xenopus embryos or microdissected anterior foregut tissue (from pharynx to stomach) were pooled and lysed in Triton X-100 lysis buffer (0.5% Triton X-100, 0.1M HEPES, 150 mM NaCl, 2mM EDTA pH 8.0, 2mM EGTA) containing protease inhibitors (ThermoFisher). Protein lysates were combined with 2X Laemmli buffer and 0.1M DTT, boiled, and separated by 4-12% Bis-Tris gel electrophoresis (ThermoFisher). For HRP detection, membranes were blocked in 5% BSA, then incubated in primary antibody (1:1000) overnight in blocking solution at 4°C. Membranes were incubated in HRP-conjugated secondary antibody (1:5000) in blocking solution for 1 hour at room temperature, then visualized by addition of ECL substrate (Amersham, GE) and imaged using a ChemiDoc (BioRad). Western blot quantification was performed in ImageLab (BioRad).

## Resources

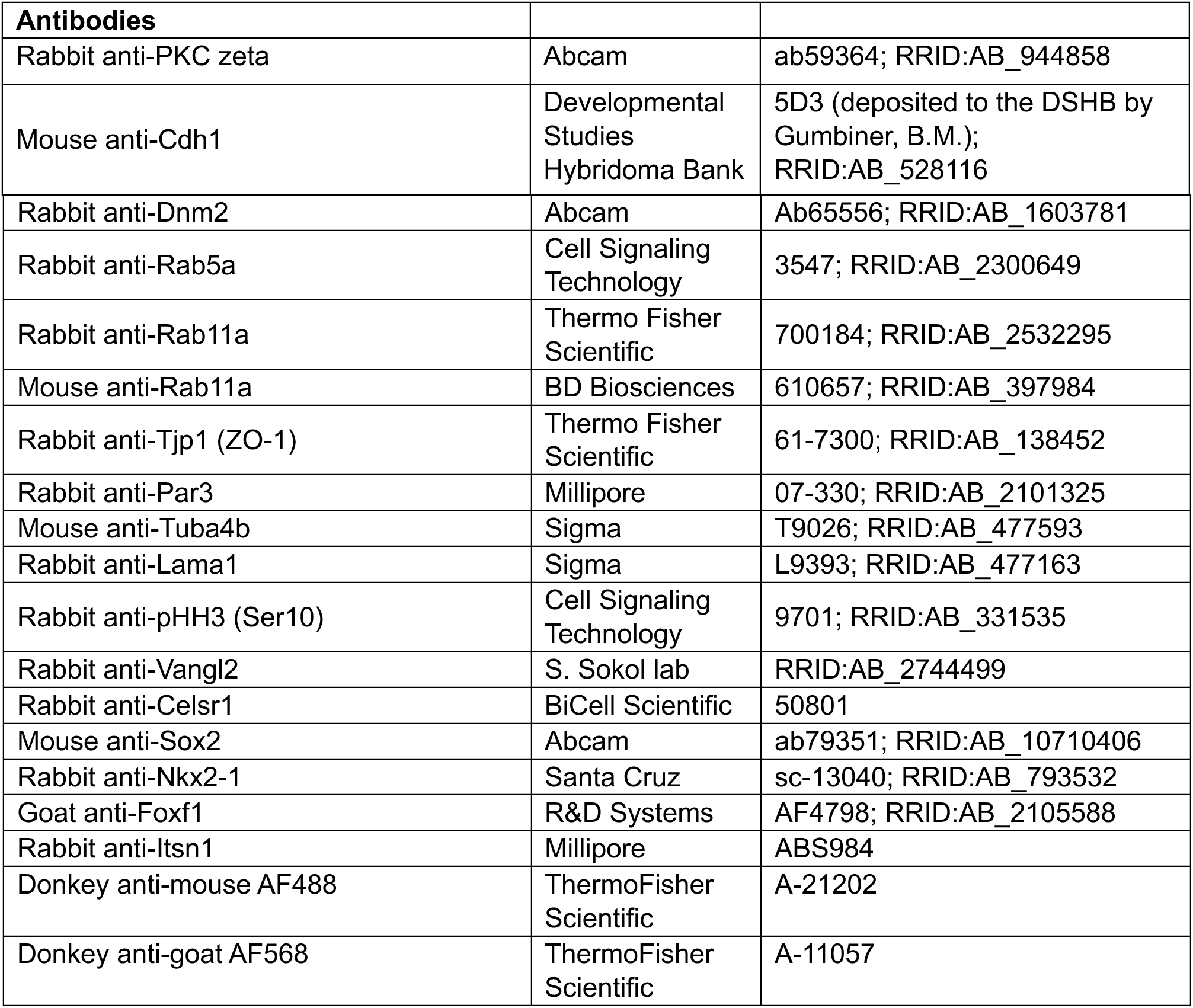
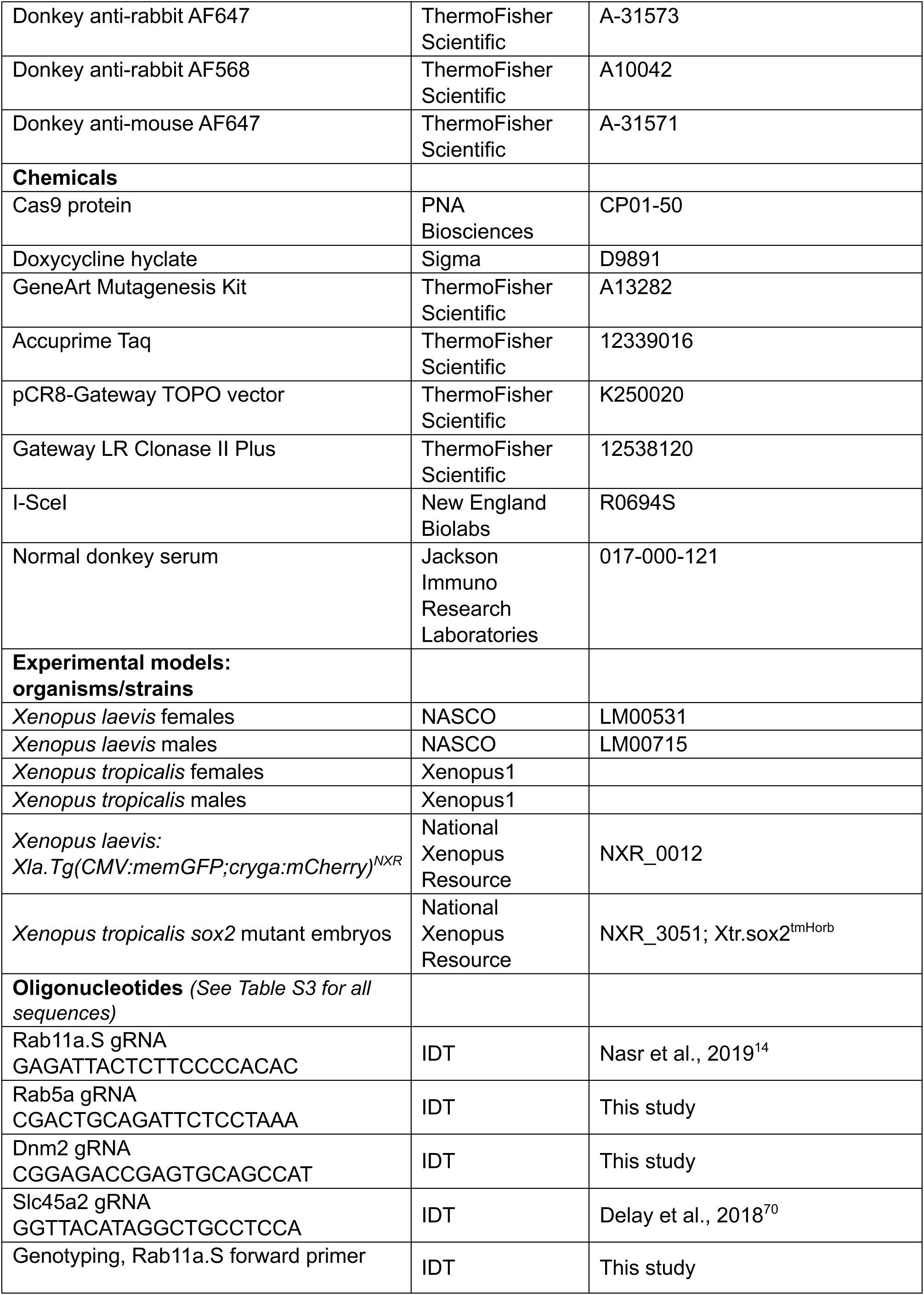
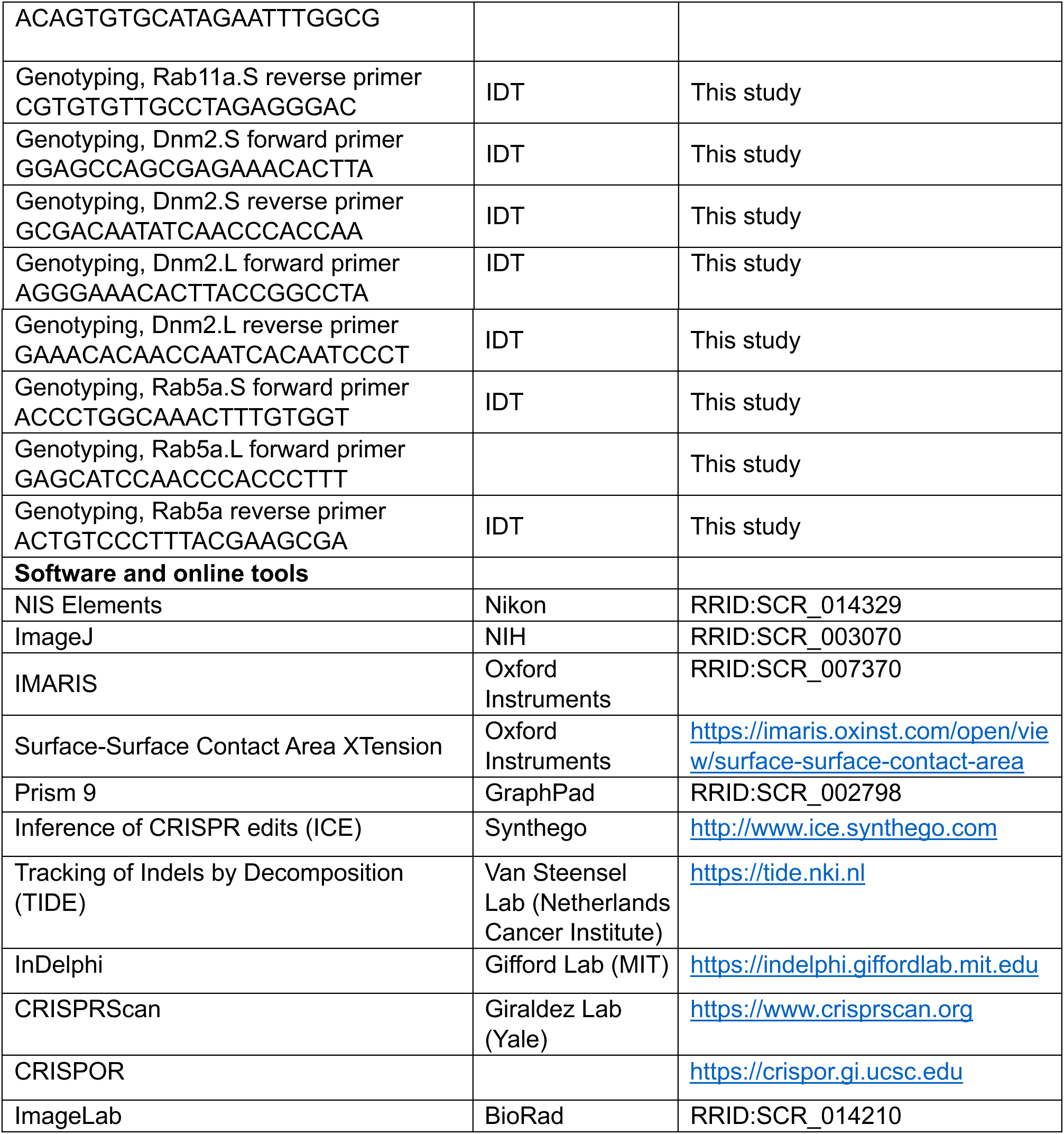

---

**Supplemental Figure 1.**
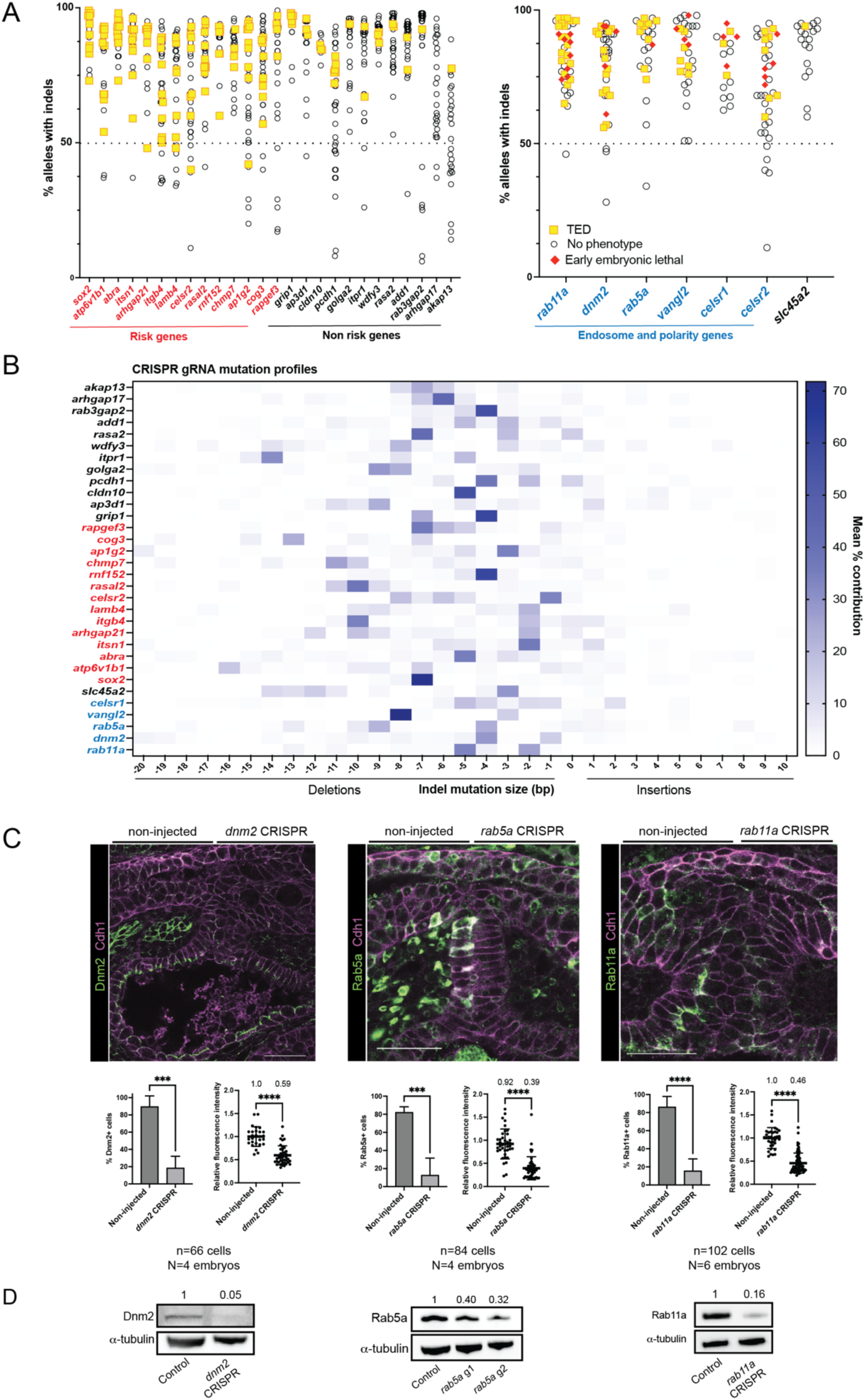
(Related to Figure 1) F0 CRISPR-Cas9 editing strategy generates highly edited embryos. A) Left: Graph demonstrating correlation between editing efficiency (% alleles with indels) and phenotype penetrance in embryos with mutations in patient risk genes. Right: Graph shows the correlation between editing efficiency, embryonic lethality, and phenotype penetrance in embryos with mutations in endosome and polarity genes. B) Heat map displaying the frequency and types of indel mutations generated by the indicated gRNA in F0 embryo, mean of n=5-15 individual embryo Sanger sequencing analysis per gRNA. C) Immunostaining and confocal microscopy of unilaterally-injected tadpoles at the 2-cell stage with dnm2, rab5a, or rab11a CRISPR reagents and quantification of reduced number of protein-expressing cells and protein abundance on the injected side of the tadpole compared to the control, non-injected side. Graphs are mean ± SEM, ***p<0.001, ****p<0.0001 Students’ t-test, numbers above dot plots are the mean. D) Immunoblotting of foreguts dissected from Xenopus CRISPR mutants compared to controls. Relative protein abundance indicated by numbers above immunoblots, with control protein abundance normalized to 1.

**Supplemental Figure 2.**
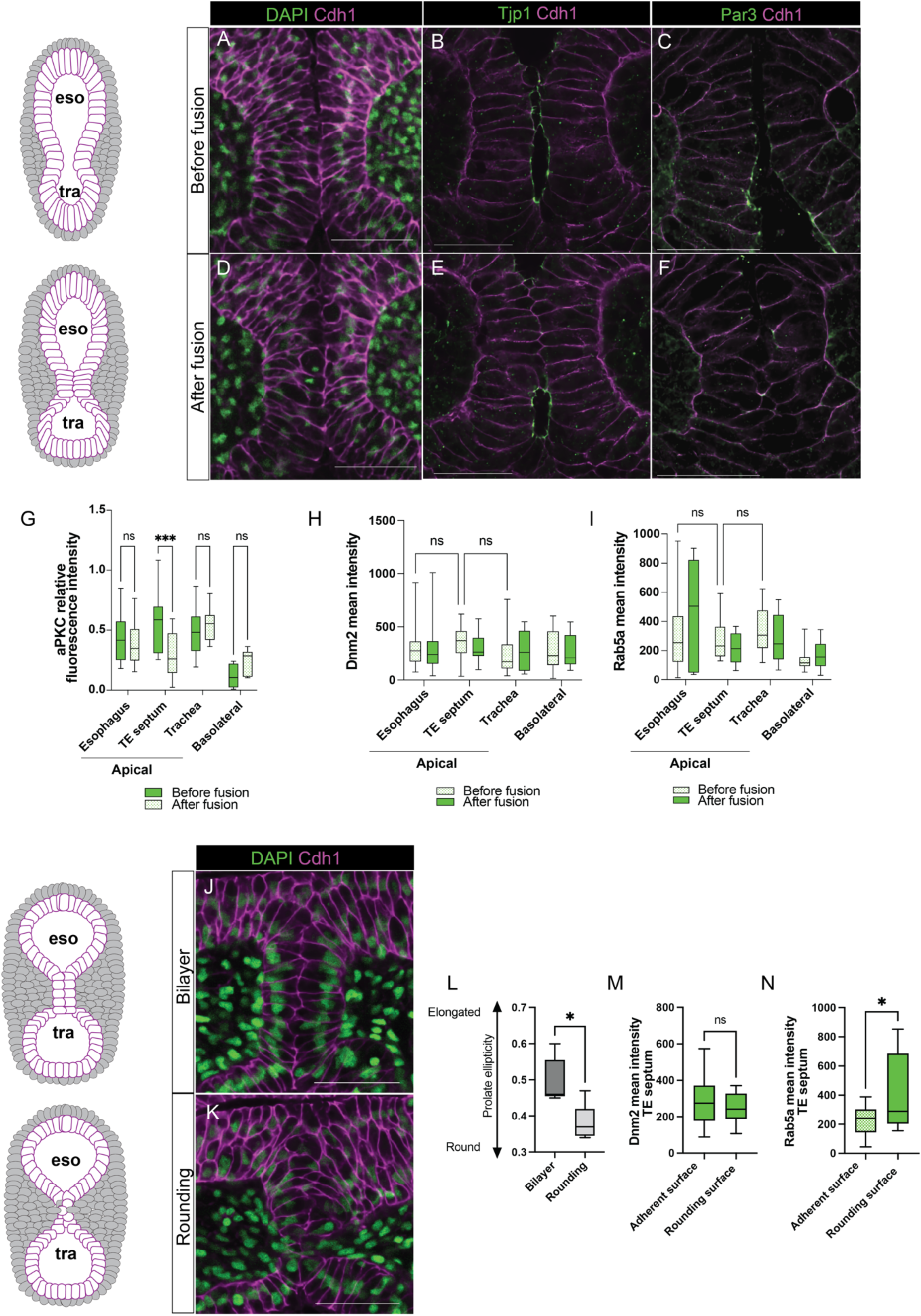
(Related to Figure 2) Dynamic cell shape and polarity changes during trachea-esophageal separation. A-C) Before fusion the foregut is a pseudostratified epithelium with apical tight junctions (Tjp1) and Par3. D-F) After apical fusion, epithelial cells remodel to localize E-Cadherin (Cdh1) apically and downregulate Tjp1 and Par3. G) Quantification of aPKC downregulation after epithelial fusion (mean ± min/max, ****p<0.00001, 2W-ANOVA, n=6 cells per embryo, N=5 embryos). H) Quantification of Dnm2 intensity after epithelial fusion (mean ± min/max, 2W-ANOVA, n=6 cells per embryo, N=5 embryos). I) Quantification of Rab5a intensity after epithelial fusion (mean ± min/max, 2W-ANOVA, n=6 cells per embryo, N=5 embryos). J) Before resolution of the septum, the foregut epithelium remodels to a bilayered columnar epithelium with basally positioned nuclei. K) The septum resolves after downregulation of E-Cadherin and integration of septum cells into either the trachea or esophagus. L) Prolate ellipticity is significantly decreased in cells actively resolving in the septum (mean ± min/max, *p<0.05, Student’s t-test, n=6 cells per embryo, N=5 embryos). M) Quantification of Dnm2 intensity before and after resolution (mean ± min/max, Student’s t-test, n=6 cells per embryo, N=5 embryos) N) Quantification of Rab5a intensity before and after resolution (mean ± min/max, *p<0.05, Student’s t-test, n=6 cells per embryo, N=5 embryos).

**Supplemental Figure 4.**
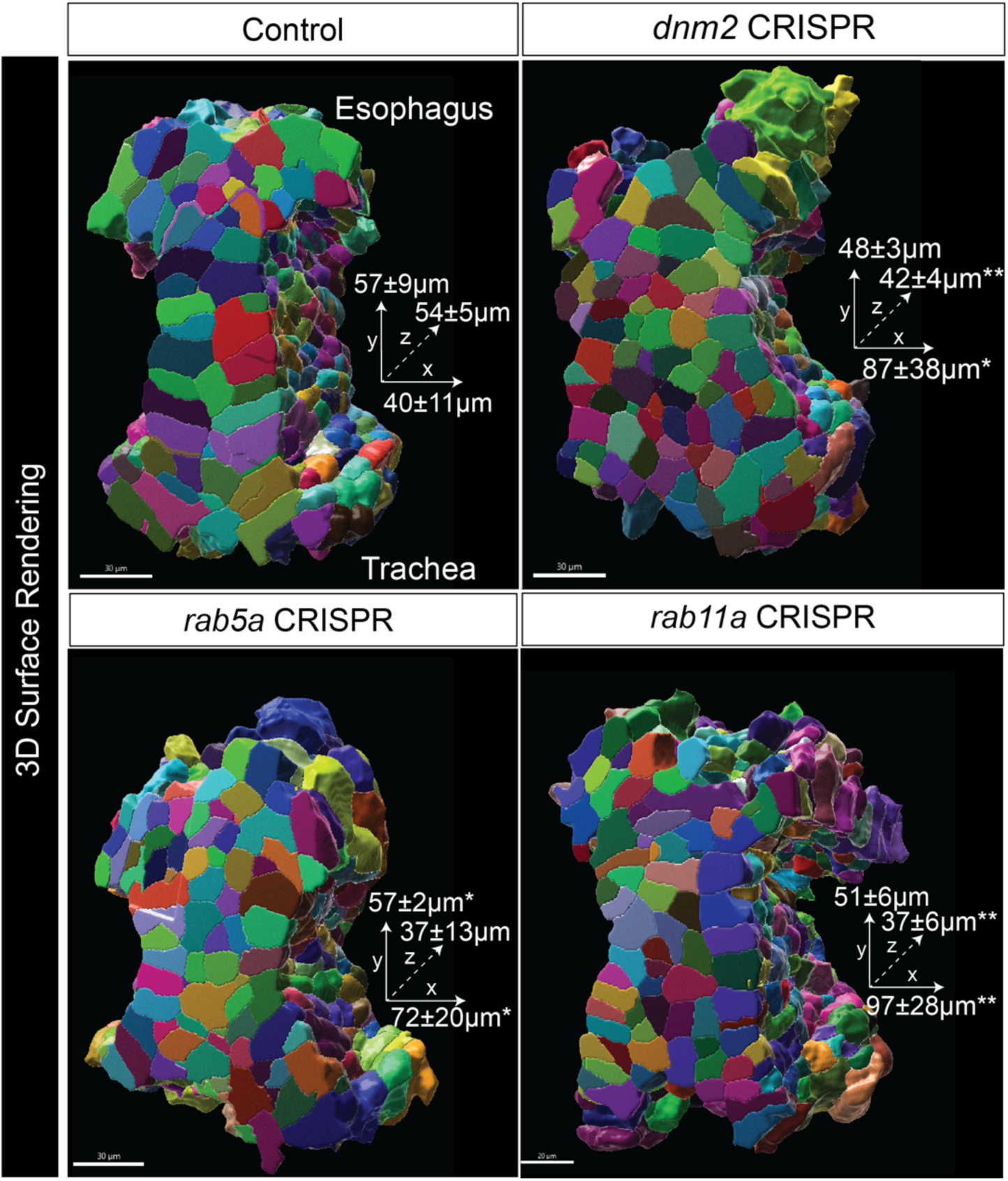
**(Related to Figure 4) 3D surface renderings** and measurements of the epithelial septum showing that mutants are significantly wider (x axis) and shorter in the rostral-caudal (z axis) axis compared to controls (mean length ± SEM, Student’s t-test compared to control lengths, *p<0.05, **p<0.01, n=4-6 embryos per condition). Scale bars are 30 µm.

**Supplemental Figure 5.**
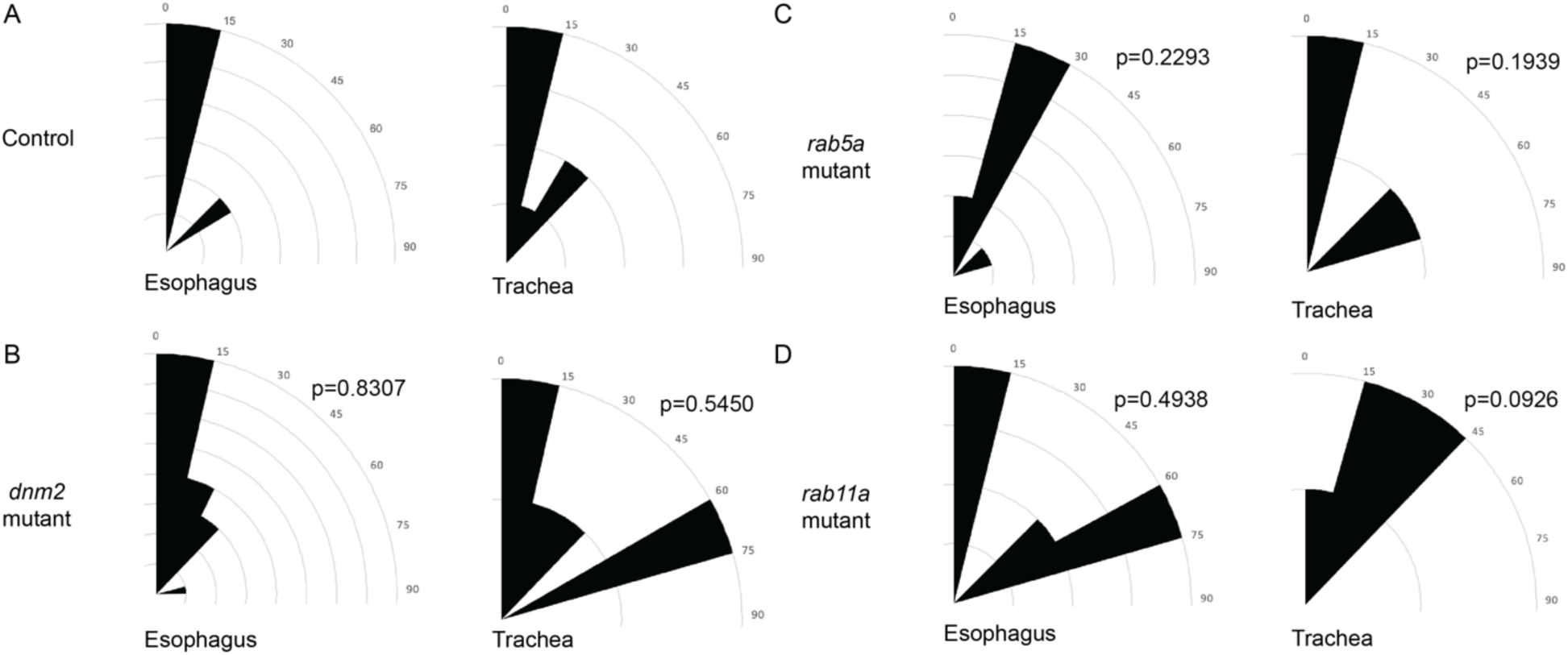
(Related to Figure 5). Mitotic spindle angles are not significantly changed compared to controls in the trachea and esophageal epithelia of endosome mutants. A) Control embryo trachea and esophagus epithelia most frequently divide between 0- 15°C relative to the plane of the tissue. B-D) Endosome mutant trachea and esophagus epithelia division angles are not significantly altered, but trend to be more variable than control trachea and esophageal epithelia (Kolmogorov-Smirnov test, n.s. p>0.05, 3-5 cells measured per embryo, 5 embryos measured per condition).

**Supplemental Figure 6.**
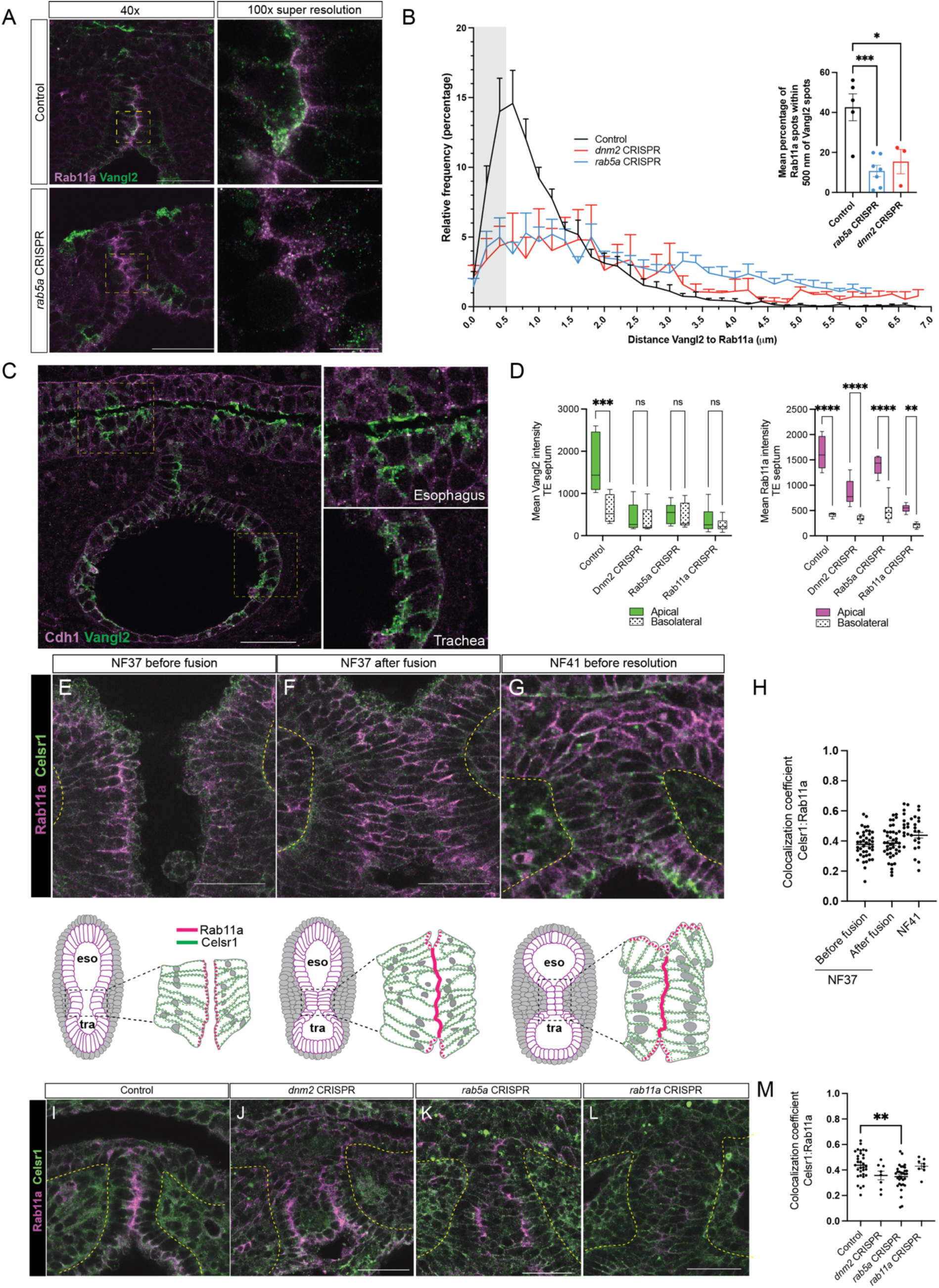
(Related to Figure 6) Celsr1 is localized to the periphery of cells during tracheoesophageal separation. A) Super-resolution confocal of Rab11a-Vangl2 immunostaining in representative control and Rab5a CRISPR mutant embryos. B) Quantification of distance between Vangl2-Rab11a spots binned and plotted as relative frequency histograms. Inset: mean percentage of Vangl2 spots within 500nm of Rab11a spots (area under curve in gray box) mean ± SEM, N=3-7 embryos, *p<0.05. ***p<0.001 C) Immunostaining of Vangl2 in the trachea and esophageal epithelium. Vangl2 is localized apically in the lumen-facing esophageal epithelia but not in the more basal cell layers and is localized around the membrane periphery in the single cell tracheal epithelium (insets). D) Vangl2 intensity at the apical membrane is decreased in dnm2, rab5a, and rab11a CRISPR mutants, while Rab11a remains localized to the apical membranes (mean ± min/max, 2W-ANOVA, ****p<0.0001, ***p<0.001, **p<0.01, n=3-5 cells per embryo, 5 embryos analyzed). E-H) Time course of Celsr1 immunostaining during *Xenopus* trachea-esophageal morphogenesis. Diagrams depict Celsr1/Rab11a localization during foregut separation. I-M) Celsr1:Rab11a colocalization is significantly decreased in *rab5a* Xenopus mutants, but Celsr1 cellular localization is not significantly altered in *dnm2*, *rab5a*, *and rab11a* Xenopus mutants (mean ± min/max, 1W-ANOVA, **p<0.01. n=3-5 cells, 5 embryos analyzed).

**Table S1.**

| ***Xenopus* ortholog of patient gene** | **Protein Function** | **% Indel mutations (mean ± SD)** | **Mean frequency of trachea-esophageal defect (±SEM, n, N, p value)** | ***Xenopus* Phenotype** |
| --- | --- | --- | --- | --- |
| *sox2* | Transcription factor | 100% -germline null | 100% (±0, 14, 2, $p < 0.0001$)*** | TEF – 14/14 |
| *sox2* | Transcription factor | 90%± 8.6 | 61% (±10, 18, 3, $p < 0.0001$)*** | TEF – 11/18 |
| ***atp6v1b1*** | Intracellular transport | 82%±11.5 | 40% (±19,30, 2, $p = 0.0024$)** | TEF – 4/30<br>Occluded lumens – 9/30 |
| *abra*<sup>^</sup> | Rho-actin regulation | 90%±6.1 | 41% (±8, 29, 4, $p = 0.0009$)** | TEF – 10/29<br>Occluded lumens – 4/29 |
| *itsn1*<sup>^</sup> | Endocytosis, Cdc42 GEF | 87%±8.2 | 42% (±13, 26, 3, $p = 0.0038$)** | TEF – 6/26<br>Occluded lumens – 5/26 |
| *arhgap21*<sup>^</sup> | Rho family GTPase | 85%±12.0 | 39% (±7, 23, 4, $p = 0.0006$)** | TEF – 5/23<br>Occluded lumens – 8/23 |
| *itgb4*<sup>^</sup> | Integrin adhesion, putative cargo | 72%±14.5 | 26% (±8, 43, 5, $p = 0.0063$)** | TEF – 9/43<br>Occluded lumens – 4/43 |
| ***lamb4*** | Laminin subunit | 71%±17.5 | 28% (±19, 32, 3, $p = 0.0121$)* | TEF - 9/32<br>Occluded lumens – 1/32 |
| *celsr2*<sup>^</sup> | Cell polarity, putative cargo | 72%±16 | 24% (±5, 33, 2, $p = 0.0041$)** | TEF - 6/33<br>Occluded lumens – 2/33 |
| ***rasal2*** | Ras GTPase-like | 81%±11.9 | 34% (±2, 32, 3, $p = 0.0005$)*** | TEF - 5/32<br>Occluded lumens – 5/32 |
| ***rnf152*** | E3 ubiquitin ligase | 85% ±11.3 | 32% (±2, 25, 3, $p = 0.0002$)*** | TEF – 7/25<br>Occluded lumens – 2/25 |
| ***chmp7*** | Late endosomes | 84%±10 | 26% (±7, 25, 3, $p = 0.0061$)** | TEF - 6/25<br>Occluded lumens - 1/25 |
| *ap1g2*<sup>^</sup> | Endocytosis | 76%±22.1 | 23% (±3, 47, 5, $p = 0.0038$)** | TEF – 7/47 |
| ***cog3*** | ER-Golgi transport | 79%±14.5 | 18% (±6, 47, 3, $p = 0.037$)* | TEF - 9/47<br>Occluded lumens - 3/47 |
| *rapgef3*<sup>^</sup> | Ras signaling | 86%±13 | 15% (±3, 34, 3, $p = 0.0360$)* | TEF – 4/34<br>Occluded lumens – 2/34 |
| ***grip1*** | Scaffold protein | 95%±4.5 | 17% (±8, 44, 3, $p = 0.0712$) | |
| ***ap3d1*** | Intracellular transport | 85%±13.8 | 15% (±7, 20, 2, $p = 0.0808$) | - |
| ***cldn10*** | Tight junctions | 78%±11.3 | 14% (±7, 34, 3, $p = 0.1106$) | - |
| *pcdh1*<sup>^</sup> | Cell adhesion, putative cargo | 72%±14 | 13% (±7,34, 3, $p = 0.1361$) | - |
| ***golga2*** | Golgi transport | 87%±10 | 12% (±3, 38, 3, $p = 0.0606$) | |
| ***itpr1*** | Inositol triphosphate receptor | 79%±17.5 | 8% (±14, 27, 3, $p = 0.4262$) | - |
| ***wdfy3*** | Autophagy | 90%±4 | 9% (±5, 31, 3, $p = 0.2680$) | |

**Table S1.**
| *rasa2* | Ras GTPase | 85%±11 | 9% (±8, 38, 4, p=0.1950) | - |
| --- | --- | --- | --- | --- |
| *add1* | Cytoskeleton | 87%±9.3 | 5% (±5, 39, 3, p=0.6608) | - |
| *rab3gap2*<sup>^</sup> | Endocytosis | 88%±14.2 | 5% (±6, 34, 3, p=0.5948) | - |
| *arhgap17*<sup>^</sup> | Rho signaling | 75%±16 | 4% (±4, 39, 5, p=0.7341) | - |
| *akap13* | Rho guanine exchange factor | 58%±14.5 | 6% (±6, 29, 3 p=0.5165) | - |
| *tyr* (control) | Pigmentation | n/d | 2% (±2, 36, 6) | - |
| Endosome/polarity genes | Protein Function | % Indel mutations (mean ± SD) | Mean frequency of trachea-esophageal defect (±SEM, n, N, p value) | Xenopus Phenotype |
| *dnm2* | Endocytosis | 82%±17 | 70% (±6, 65, 16, p=0.0002)*** | TEF – 45/65<br>Occluded lumens – 14/65 |
| *rab5a* | Early endosome GTPase | 79%±9 | 63% (±6, 78, 16, p=0.0004)*** | TEF – 41/78<br>Occluded lumens – 9/78 |
| *rab11a* | Recycling endosome GTPase | 86%±18 | 60% (±7, 46, 18, p=0.0039)** | TEF – 23/46<br>Occluded lumens – 5/46 |
| *vangl2* | Cell polarity, putative cargo | 69%±18 | 34% (±14, 25, 3, p=0.0141)* | TEF – 7/25<br>Occluded lumens – 1/25 |
| *celsr1* | Cell polarity, putative cargo | 80%±10 | 33% (±4, 20, 3, p=0.0008)*** | TEF - 6/20<br>Occluded lumens – 1/20 |
| *slc45a2* (control) | Pigmentation | 82%±14.8 | 2% (±3, 26, 3) | - |

<sup>^</sup>Genes reported in Zhong et al., HGG Advances 2022 (n=10)
***New genes reported in this study (n=15)***
TEF – trachea-esophageal fistula
SEM – standard error of the mean, n=number of embryos assayed, N=number of independent experiments p value determined by unpaired Student's t-test compared to *tyr* controls. p<0.05 indicates a significant frequency of TEF over controls.

